# Amphetamine and Nicotine Reduce Sucrose Self-Administration Independent of Sex

**DOI:** 10.1101/2025.06.20.660788

**Authors:** Cameron Fulco, Carlos Cordero, Katelyn George, Aaron G Roseberry

## Abstract

Amphetamine and nicotine are two widely used and abused drugs that are taken for legitimate pharmaceutical purposes but are also highly abused through illicit recreational use. Both of these drugs have been widely shown to decrease food intake in both humans and pre-clinical models, and although amphetamine and nicotine clearly affect food intake under normal baseline (‘homeostatic’) conditions, there has been limited examination of the ability of these drugs to affect reward-related (‘hedonic’) aspects of feeding. Furthermore, there are sex differences in the behavioral responses to both drugs, but it is unclear if these sex differences also translate to their effects on feeding. This study examined whether nicotine and amphetamine regulate sucrose intake in a food self-administration paradigm in a sex-dependent manner across both fixed and progressive schedules of reinforcement. Amphetamine reduced operant responding for sucrose pellets and decreased acute intake of sucrose during *ad libitum* free-feeding access in a dose-dependent manner, whereas nicotine reduced sucrose self-administration and free intake only at higher doses that also impaired locomotor activity in open field tests. The effects of both amphetamine and nicotine did not differ by sex for either drug. Overall, these results suggest that the mechanisms mediating the addictive qualities of these drugs and their appetite suppressing effects may be distinct and therefore could be a potential target for future obesity therapeutics.

## Introduction

Worldwide obesity rates have significantly increased over the past few decades, tripling in prevalence since 1974 and currently affecting 13% of the total population (World Health Organization, 2018). While the increased weight associated with obesity is a key factor in a shortened lifespan, it also exacerbates other health problems such as diabetes mellitus, heart disease, cancers, osteoarthritis and sleeping disorders which highlights the need for development of therapeutic interventions (Kinlen et al., 2018; Ogunbode et al., 2011). The consistent rise in the prevalence of obesity has largely been attributed to maladaptive alterations in feeding (Aronne et al., 2009), which is a complex behavior that is regulated by multiple peripheral and central molecules and circuits.

The mesolimbic dopamine system is the primary neural circuit mediating reward, reinforcement and motivated behaviors, and it is heavily involved in multiple aspects of feeding (Fallon et al., 2007; Volkow & Wise, 2005; Wise, 2006; Ungerstedt, 1971; Cannon et al., 2004a, 2004b). Dopamine neurons in the ventral tegmental area (VTA) form the center of the mesolimbic dopamine system and project to multiple efferent targets including the nucleus accumbens, amygdala, hippocampus, olfactory tubercle and others (for review, see Morales & Margolis, 2017), and dopamine has been shown to be increased in many of these areas in response to food and food related cues (Fallon et al., 2007; Fuchs et al., 2005). Dopamine also modulates food seeking behaviors (Roitman et al., 2004) and enhances the incentive salience of palatable foods (DiFeliceantonio & Berridge, 2016). Finally, it appears that dopamine must remain at an optimal level for feeding to occur, as both dopamine depletion and increased dopamine release inhibit feeding (Heffner et al., 1977; Ungerstedt, 1971, Zhou & Palmiter, 1995).

Dopamine circuits are a major target for abused drugs which all increase dopamine release and induce addictive behaviors due to their activation of the mesolimbic dopamine system (Grace, 2000). Interestingly, many abused drugs also show strong effects on feeding (Kraeuchi et al., 1985; Balopole et al., 1979; Morley & Flood, 1987; Wellman et al., 2002). Thus, identifying how abused drugs inhibit feeding independent of their rewarding, reinforcing, and addictive qualities may allow for new strategies to combat excessive feeding and obesity. Amphetamine and nicotine are two commonly used drugs of abuse which act on the mesolimbic dopamine system and show strong effects on feeding. While both of these drugs act on the dopamine system, their mechanisms of action differ.Amphetamine primary acts on the dopamine system by blocking dopamine transporters and causing reverse efflux of dopamine through them to increase extracellular dopamine levels (Fleckenstein et al., 2007) whereas nicotine acts on the dopamine system by directly increasing dopamine neuron activity via β2-containing nicotinic acetylcholine receptors and increased excitatory drive onto dopamine neurons (Wittenberg et al., 2020).

Amphetamine was one of the first drugs used as an appetite suppressant to treat obesity. Due to its harmful side effects and addictive potential, it was removed as a treatment option, however (Stăcescu et al., 2020). Although the exact mechanisms by which amphetamine decreases feeding are unclear, they have been suggested to result from increased striatal dopamine release and subsequent inhibitory action of the orexigenic neuropeptide, Neuropeptide Y (NPY) (Li & Pelletier, 1986; Obuchowicz, 1996; Pelletier & Simard, 1991, Cannon et al., 2004a, 2004b). Although amphetamine has been widely shown to inhibit feeding under normal *ad libitum* feeding conditions, less is known about whether it affects the rewarding or motivational qualities of food. We have previously shown that amphetamine has a dose-dependent, bidirectional effect on two binge-like models of feeding (West et al., 2019), whereby lower doses of amphetamine increased sucrose intake while higher doses showed the typical decrease in intake. Although these studies were designed to test effects on the rewarding aspects of feeding, it is unclear if changes in basal consummatory behavior could have contributed to these responses, however.

Nicotine is another widely abused drug that both acts on dopamine circuits and has been widely shown to decrease food intake (Iravani, 1972; Passey et al., 1961; Schechter & Cook, 1976). Unlike amphetamine however, nicotine has been reported to affect feeding behaviors via multiple mechanisms including the regulation of CART (Dandekar et al., 2011), NPY (Chen et al., 2007; Frankish et al., 1995), melanocortin pathways (Mineur et al., 2011), endocannabinoids (Bura et al., 2010), CRH (Kamdi et al., 2009), and dopamine (Grady et al., 2007; De Biasi & Dani, 2011). As with amphetamine, nicotine’s effects on feeding under normal *ad libitum* feeding conditions has been broadly studied, but its potential effects on the rewarding and motivational qualities of food are unknown.

Sex differences have also been widely identified in both the control of feeding, and body weight, and the responses to drugs of abuse (for review, see Shi & Clegg, 2009; Becker & Hu, 2008). For example, males and females differ in basal energy expenditure and metabolic processes (Morio et al., 1997; Carpenter et al., 1998, Romanski et al., 2000; Hellström et al., 1996; Leibel & Hirsch, 1987), as well as peripheral and central mechanisms controlling feeding (Geary et al., 1994; Clegg et al., 2007; Rosenbaum et al., 1996; Munro et al., 2006; Sinclair et al., 2017). Humans and animal models also exhibit clear differences in the responses to drugs of abuse, with females showing higher levels of drug self-administration and drug intake, increased rates of drug acquisition, increased locomotor responses to drugs, higher levels of intake, and increased drug craving during abstinence compared to males (Zakiniaeiz et al., 2019, Milesi-Hallé et al., 2007, Shahbazi et al., 2008, Donny et al., 2000, Lynch, 2009, Leventhal et al., 2007, Wang et al., 2014; Locklear et al., 2012) for a variety of drugs including both amphetamine and nicotine. It is unclear whether the sex differences in these responses to drugs extends to sex differences in their effects on feeding, however. In these studies, we have used sucrose self-administration in both operant chambers and the home cage to test whether amphetamine and nicotine affect sucrose intake under more motivational and rewarding conditions and whether there were sex differences in the effects of these drugs on sucrose intake in these paradigms.

## Materials & Methods

### Animals

Young adult male and female C57Bl/6J mice (Jackson Laboratories, Bar Harbor, Maine) were used for all experiments. Mice were ten weeks of age at the start of all experiments. Mice were housed on a 12:12 light:dark cycle with *ad libitum* access to standard chow (3.36 kcal/g, Rodent Laboratory Chow 5001) and water throughout the study, and all experiments were conducted at the beginning of the dark phase. All protocols and procedures were approved by the Institutional Animal Care and Use Committee at Georgia State University and conformed to the National Research Council *Guide for the Care and Use of Laboratory Animals*.

### Reagents

Sterile bacteriostatic saline was from Patterson Veterinary Supply (Loveland, CO). d-amphetamine was from Sigma-Aldrich (St. Louis, MO). Nicotine tartrate was from the National Institute on Drug Abuse Drug Supply Program. The sucrose pellets were from Bio-Serv (Flemington, NJ). FED devices were made in-house using electronics purchased from Open Ephys (Lisbon, Portugal). All other reagents were from common commercial sources.

### Experimental Approach

Nicotine tartrate and d-Amphetamine were dissolved in sterile saline, which served as control injections for both drugs. Nicotine doses listed are calculated as the concentrations for the nicotine tartrate salt. These doses correspond to the free doses of nicotine as follows (0.1 mg/kg = 35μg/kg, 0.5 mg/kg = 175μg/kg, 1 mg/kg = 0.35mg/kg, 2 mg/kg = 0.7mg/kg, 5 mg/kg = 1.75mg/kg). Injections (10 microliter/g body weight i.p.) were done at the onset of the dark phase (∼0 – 1 hours into the start of the dark phase) for all experiments. For experiments using the FED3/FED3.1 devices in the home cage, mice had continuous access to the FED devices in their home cage for 1 week prior to the onset of testing. During testing, mice had 5 days exposure to the FED devices followed by 2 days of *ad libitum* access to normal chow. FED devices were replaced at the same time daily. For the operant chamber self-administration experiments, mice underwent self-administration daily for 5 days followed by no testing for 2 days. Mice had at least one day access to the FED devices or the operant chambers prior to test days, which were separated by a minimum of 2 days. For all experiments, the order of injections was counterbalanced so that two different experimental doses were delivered to half of the mice each test day. Due to counterbalancing, half of the mice received the doses in a different order. The experimenter was blind to the doses during treatment but was unblinded for data analysis. Mice were tested in multiple cohorts, with both males and females included in each cohort.

### Operant Chamber Sucrose Self Administration

Operant chamber sucrose self-administration experiments were done in operant test boxes enclosed in sound-attenuating cubicles (Med Associates, Fairfax, Virginia). Each test box contained two nose poke ports with a food receptacle between them. During trials, both nose poke port lights, the house light, and a 2900 Hz tone were on. Entry into the active port resulted in the delivery of one 20mg sucrose pellet to the receptable port and activation of the receptacle port light, whereas activation of the inactive port had no consequence. Pellet delivery resulted in a 20 second timeout period, where the house light, tone, and nose poke port lights turned off, while the food receptacle light remained on until a head entry into the port was detected. The house light, tone, and nose poke port lights remained off for the full 20 seconds of the time out period. Nose poke entries into the active port during the time out period were recorded, but did not have any consequence. Mice underwent 60-minute sessions 5 days/week (M-F) and had free access to chow and water in their home cage between sessions. The total number of pellets remaining in the test box at the end of the session was recorded to allow for accurate measurement of the total number of pellets obtained and eaten.

### Fixed Ratio Self-Administration

At the onset of self-administration training mice first received a single 1-hour magazine training session. During magazine training mice were placed into the test chambers with cues and timeout mimicking that of the SA trials. However, mice did not need to poke for delivery of a sucrose pellet; instead, one pellet was delivered automatically every two minutes for the duration of the trial. The mice were then shifted to a fixed ratio (FR) 1 schedule, where one poke of the active port resulted in the delivery of one pellet, until a 3:1 active port to inactive port interaction was reached for three consecutive sessions.

Upon reaching the FR1 criteria, mice were moved to a FR3 schedule, where three pokes of the active port resulted in the delivery of one pellet until stable responding was obtained at this ratio (< 20% variance of eaten pellets over three days). Mice then underwent experimental testing. Test days were separated by 3-4 days (i.e. Tuesday and Friday) and mice continued FR3 sessions on non-test days (M-F).

### Progressive ratio Self-Administration

For progressive ratio trials, mice underwent the same magazine and FR1 training described above. After reaching stable responding on FR1, mice were moved to a FR3 schedule for two days. Subsequently, training changed to a progressive ratio (PR) schedule where pellet deliveries were determined by a geometric progression (*n_j_* = *5e^j/5^*-*5*) (Richardson and Roberts, 1996). Mice underwent PR training trials until stable responding was obtained at this ratio (< 20% variance of eaten pellets over three days). Once this criterion was reached for the PR schedule, mice underwent experimental testing as described above.

### Home Cage Testing

Feeding Experiment Devices, version 3 and 3.1 (FED3) were used for all home cage feeding experiments. In free-feeding conditions, FED3 devices delivered one 20mg sucrose pellet whenever the food port was empty, which allowed for normal *ad libitum* access to food. During acclimation to the FED3 devices, home-cage chow was replaced with *ad libitum* access to sucrose pellets via the FED3 devices in the free feeding mode. For acclimation mice had 2 days of sucrose/FED3 access, followed by 1 day of home-cage chow, another 2 days of sucrose/FED3 access, and 2 further days on chow, before shifting to the testing phase. During testing, mice were provided FED3s for 5 consecutive days, with the devices replaced daily at the same time, followed by 2 days of *ad libitum* access to chow. A separate cohort of mice was tested under FR3 conditions using FED3 devices in the home cage. For FR testing, mice underwent the same acclimation procedure, followed by one week on an FR1 schedule and were then moved to an FR3 until stable responding was obtained. Upon stabilization on the FR3 schedule, mice underwent experimental testing as described above.

### Locomotor Activity

Mice were tested in an open field chamber to measure locomotor activity. The chambers (30 cm x 30 cm x 30 cm) were custom built from acrylic with PVC foam walls and floor. Mice were placed into the center of the chamber and Noldus Ethovision XT software was used to track the total distance traveled by the mice. Mice received one hour access to the chamber on day one to acclimate to the testing chamber before beginning experimental trials the following day. Each test day mice were placed into the open field chamber for 15 minutes before being removed and injected with saline or drug. Ten minutes later they were returned to the open field chamber for a one-hour trial where the total distance traveled was recorded. Injections were given daily in counterbalanced order.

### Vaginal Lavage

Vaginal lavage was performed to measure estrus cycle in a subset of the female mice. Vaginal cells were collected once a day for three days during the experimental phase for some of the female mice. Vaginal cells were flushed by administering ∼100 μL distilled water through a pipette and placing a few drops of the cell suspension on a glass slide for observation. The droplets were allowed to dry before observation via microscope. Leucocytes, cornified epithelia cells and nucleated epithelial cells were all identified and compared to estrus stage identified in McLean et al. (2012) for determination of estrus stage.

### Data Analysis and Statistics

All data are presented as mean ± SEM. Experiments were conducted using a within-subjects design so that each mouse received all treatments. Data was compiled using Microsoft Excel, analyzed using Python, and statistical analyses were performed using R. Due to non-normality and violations of homogeneity of variance within the data, generalized additive models (GAMs) were used for analysis of some experiments though the mcgv package. Drug dose, time, baseline intake group, and sex were used as independent variables depending on the best fit for the model and Huber-White standard errors were applied when heteroscedasticity was detected in the residuals of the chosen model. Post hoc comparisons were conducted using the emmeans package with Bonferroni correction applied for multiple comparisons. A significance level was set at p< 0.05, *a priori* for data analysis. For analysis examining potential differences in the response to amphetamine in mice showing low vs high baseline sucrose responding, mice were divided into two groups based on a median split of their sucrose intake after saline injections for each individual experiment for data analysis as described in Sills and Vaccarino (1996). Mice with sucrose intake higher than the median intake were placed into the high baseline intake group (HBI) and mice with sucrose intake equal to or less than the median intake were placed into the low baseline intake group (LBI).

Data from home cage testing was collected for 24-hours post drug injection on test days, data was only analyzed for 4 hours post injection for amphetamine and 2 hours post injection for nicotine. This is due to the short half-life of both of these drugs in mice, about 6-7 minutes for nicotine and about 1 hour for amphetamine, therefore these time windows were selected to capture the expected onset and duration of the effects of these drugs (Matta et al., 2007; Hicks et al., 1980).

## Results

### Operant Chamber Sucrose Self-Administration

Amphetamine and nicotine both have been widely shown to acutely inhibit feeding under normal baseline *ad libitum* feeding conditions (Cannon et al., 2004; Chen et al., 2001; Winn et al., 1982), but it is unclear whether they also affect the rewarding or motivational aspects of food. Thus, we tested whether amphetamine or nicotine affect sucrose self-administration in a FR3 schedule of reinforcement. Amphetamine dose-dependently decreased sucrose self-administration in these experiments with no differences between males and females (Figure 1). Analysis of total sucrose pellets obtained and eaten reveled significant main effects of Dose (F(4,64)=59.941, p<0.001) and Sex (F(1,16)=6.826, p<0.05), but no significant Dose x Sex interaction (F(4,64)=0.813, p=0.521). *Post hoc* analysis showed that the 1, 2, and 5 mg/kg doses significantly reduced sucrose self-administration compared to a saline control (Figure 1A). Similar results were observed when measuring active lever presses. Analysis of inactive nose pokes revealed an effect of dose (F(4,64)=7.647, p<0.001), but because the rate of inactive nose poking was very low across all trials, averaging approximately 1.7 responses per trial, it is unlikely that this effect is physiologically significant (Table 1).

**Figure 1.**
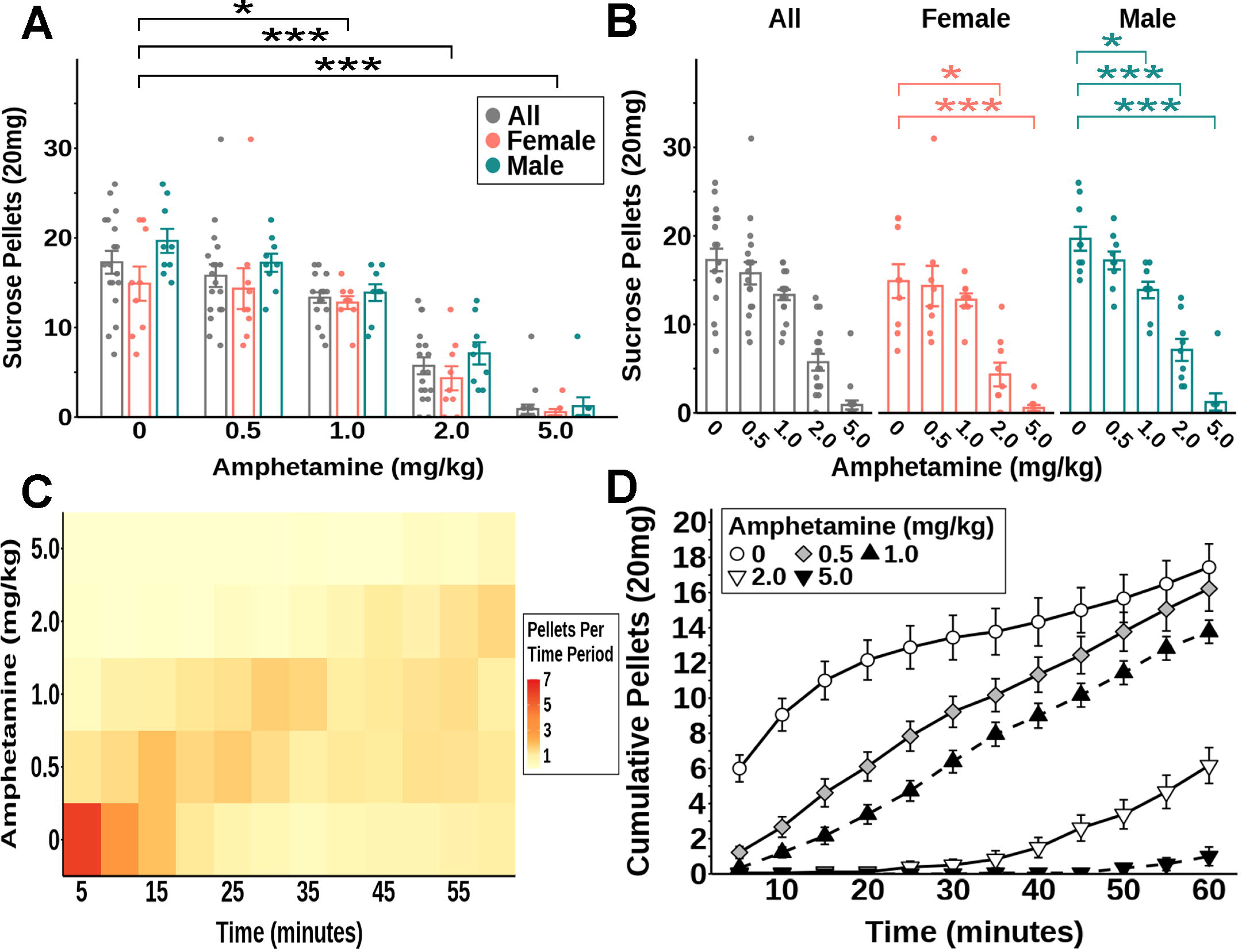
Amphetamine significantly reduced sucrose self-administration on an FR3 schedule. (**A-B**) Total self-administered sucrose pellets. (**C**) Heatmap of self-administered sucrose pellets per 5-minute interval. (**D**) Cumulative self-administered sucrose pellets over time. All: n = 18; males: n = 9; females: n = 9. * P<0.05 vs saline; ** P<0.01 vs saline; *** P<0.001 vs saline.

**Table 1.** Mean Inactive Lever Presses Per Experiment

| **Amphetamine** |  |  |  |  |  |  |  |
| --- | --- | --- | --- | --- | --- | --- | --- |
|  | Experiment | Saline |  | 0.5 mg/kg | 1 mg/kg | 2 mg/kg | 5 mg/kg |
|  | Operant Conditioning Trails FR3 | 3.67 |  | 2.50 | 1.28 | 0.67 | 0.33 |
|  | Operant Conditioning Trails PR | 4.10 |  | 1.30 | 1.35 | 0.50 | 0.10 |
| **Nicotine** |  |  |  |  |  |  |  |
|  | Experiment | Saline | 0.1 mg/kg | 0.5 mg/kg | 1 mg/kg | 2 mg/kg | 5 mg/kg |
|  | Operant Conditioning Trails FR3 | 2.67 | 2.89 | 1.94 | 2.78 | 1.22 | 1.67 |
|  | Operant Conditioning Trails PR | 4.28 | 2.94 | 3.33 | 1.78 | 0.61 | 0.50 |

Because we were interested in whether there were sex differences in the effects of amphetamine, we also analyzed the male and female data independently even though there was not a significant dose x sex interaction. When we analyzed the male and female data independently there was a significant effect of dose for both males (F(4,32)=47.174, p<0.001) and females (F(4,32)=20.506, p<0.001). *Post hoc* analysis revealed that for males the 1, 2 and 5 mg/kg doses significantly reduced sucrose self-administration compared to a saline control and for females this was only significant for the 2 and 5 mg/kg doses (Figure 1B).

We next tested whether the baseline level of sucrose self-administration influenced the ability of amphetamine to inhibit responding because past studies have shown that baseline food intake can act as a cofactor in feeding responses to amphetamine (Sills & Vaccarino, 1996; Sills et al., 1998), an effect which was also seen if the animals were food restricted to change baseline intake (Samson, 1986). To test this, a median split of the mice was applied to the data depending on sucrose intake during the saline condition (Sills and Vaccarino, 1996). Analysis of sucrose intake including the baseline sucrose intake grouping variable revealed a main effect of dose (F(4,56)=67.654, p<0.001), baseline intake group (F(1,14)=7.991, p<0.05), and Sex (F(1,14)=9.8653, p<0.01), and a significant baseline intake group x dose interaction (F(4,56)=4.065, p<0.01). Post hoc analysis revealed that for the high baseline intake group, 1, 2 and 5 mg/kg amphetamine significantly reduced sucrose self-administration compared to control (Figure 2A), but for the low baseline intake group, only the 2 and 5 mg/kg doses significantly reduced sucrose self-administration compared to control (Figure 2A). Interestingly the high and low baseline intake groups appeared to have different dose responses to amphetamine as the high intake group was significantly inhibited by 1 mg/kg and showed a trend toward inhibition by 0.5 mg/kg amphetamine, the low baseline group had no response to either dose (Figure 2A).

**Figure 2.**
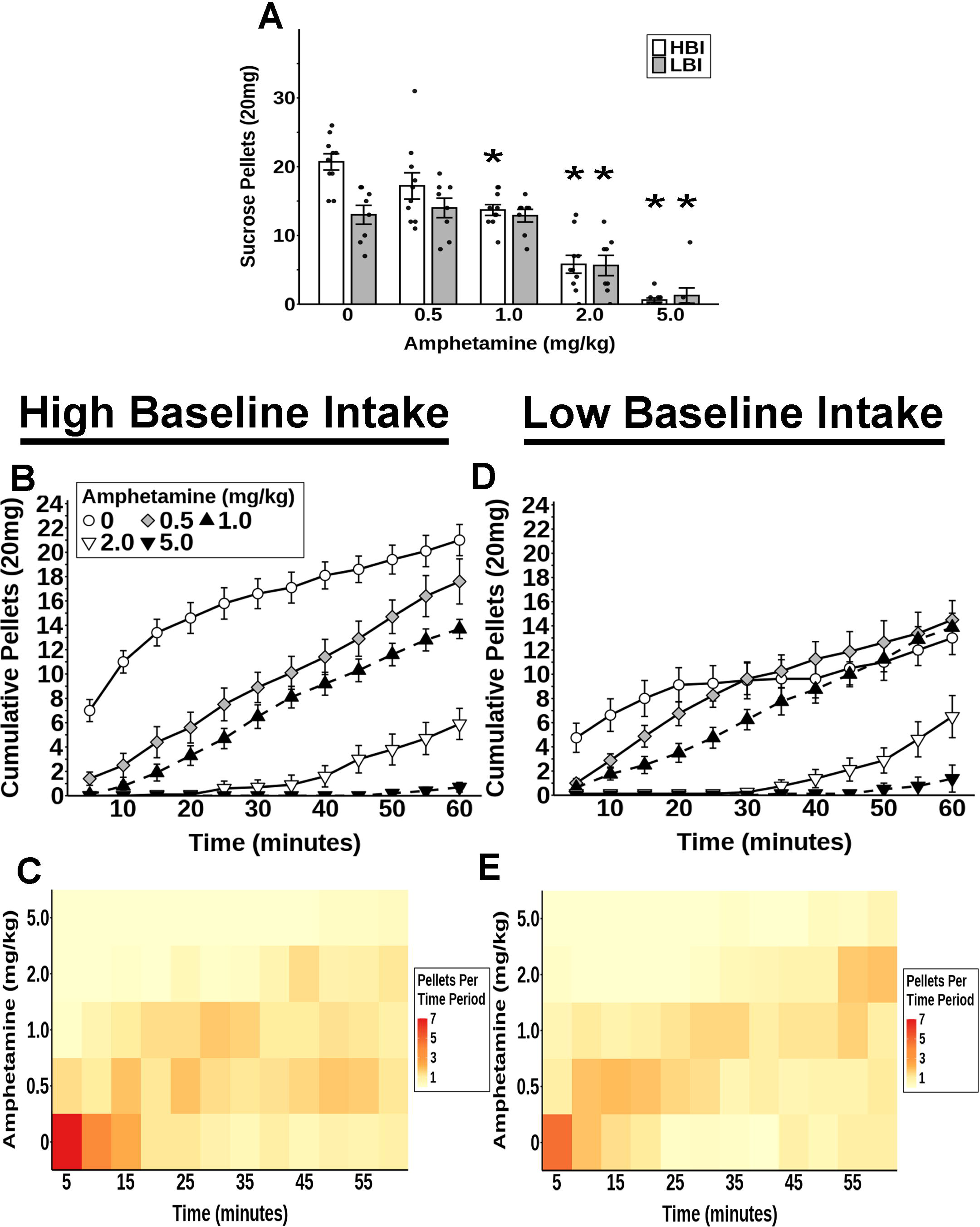
Amphetamine differentially reduced sucrose self-administration on an FR3 schedule in mice with high vs low baseline sucrose intake. (**A**) Total self-administered sucrose pellets for high and low baseline intake groups. (**B,D**) Cumulative self-administered sucrose pellets over time for mice in the high (**B**) and low (**D**) baseline sucrose intake groups. (**C,E**) Heatmap of self-administered sucrose pellets per 5-minute interval for mice in the high (**C**) and low (**E**) baseline sucrose intake groups. High Baseline Intake group: n = 10 (n = 5 males, n = 5 females). Low Baseline Intake group: n = 8 (n = 4 males, n = 4 females). * P<0.05 vs saline.

We also tested the effects of nicotine on sucrose self-administration in the same paradigm. Although analysis of sucrose intake after nicotine injections revealed a significant main effect of dose (F(5,80)=7.534, p<0.001) and a significant effect of sex (F(1,16)=4.837, p<0.05), *post hoc* analysis revealed that only the highest dose, 5 mg/kg, significantly reduced sucrose self-administration for all mice compared to control (Figure 3A). As with amphetamine, there were no sex differences in the effects of nicotine on sucrose self-administration (i.e. no dose x sex interaction: F(1,80)=1.022, p=0.410). Furthermore, nicotine did not have an effect on inactive nose pokes (F(5,80=1.525, p=0.191)). As with amphetamine, when we separated the data by sex, there was a significant effect of dose for both males (F(5,45)=5.064, p<0.001) and females (F(5,35)=3.8334, p<0.01). In contrast to amphetamine, however, *post hoc* analysis revealed that none of the doses significantly reduced sucrose self-administration for males or females (Figure 2B).

**Figure 3.**
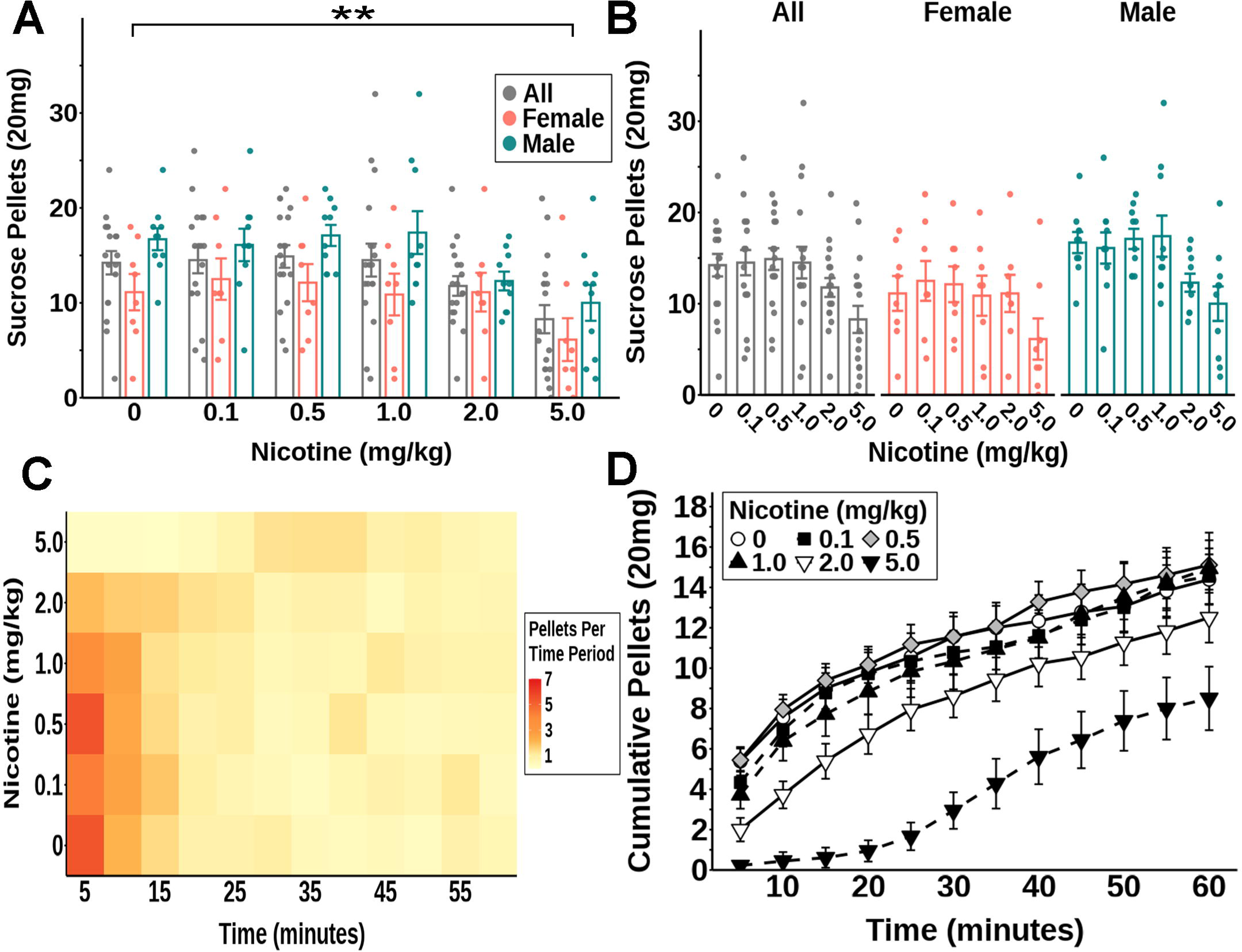
High dose nicotine significantly reduced sucrose self-administration on an FR3 schedule. (**A-B**) Total self-administered sucrose pellets. (**C**) Heatmap of self-administered sucrose pellets per 5-minute interval. (**D**) Cumulative self-administered sucrose pellets over time. All: n = 18; males: n = 10; females: n = 8. * P<0.05 vs saline; ** P<0.01 vs saline; *** P<0.001 vs saline.

Next, we tested whether amphetamine or nicotine affected the motivational qualities of sucrose pellets using a progressive ratio schedule of reinforcement. As with the FR3 data, we also tested whether the level of baseline responding affected the response to amphetamine (Figure 4E, Figure 4F). GAM analysis revealed a sex x dose interaction for amphetamine. *Post hoc* analysis revealed that 2 and 5 mg/kg doses reduced nose pokes and breakpoints compared to saline for both sexes (Figure 4B, Figure 4D), and the 2 mg/kg dose significantly reduced nose pokes and breakpoint more in females compared to males (Figure 4A, Figure 4C). Analysis of inactive nose pokes revealed a significant main effect of dose as well, but as with the FR data, because the rate of inactive nose poking was quite low across all trials, averaging approximately 1.5 responses per trial, it is unlikely that this effect is physiologically significant (Table 1). When mice were grouped based on their baseline intake, there was also a significant baseline intake x dose interaction for both breakpoint and active nose pokes. *Post hoc* analysis revealed that for the high baseline group, all doses (0.5, 1, 2, and 5 mg/kg) significantly reduced nose pokes compared to saline (Figure 4E),but only the 2 and 5 mg/kg doses significantly reduced nose pokes in the low baseline group (Figure 4E). Similar results were observed when breakpoints were analyzed with all doses significantly reducing breakpoints in the high baseline group but only the 2 and 5 mg/kg doses reducing breakpoints in the low baseline group (Figure 4F).

**Figure 4.**
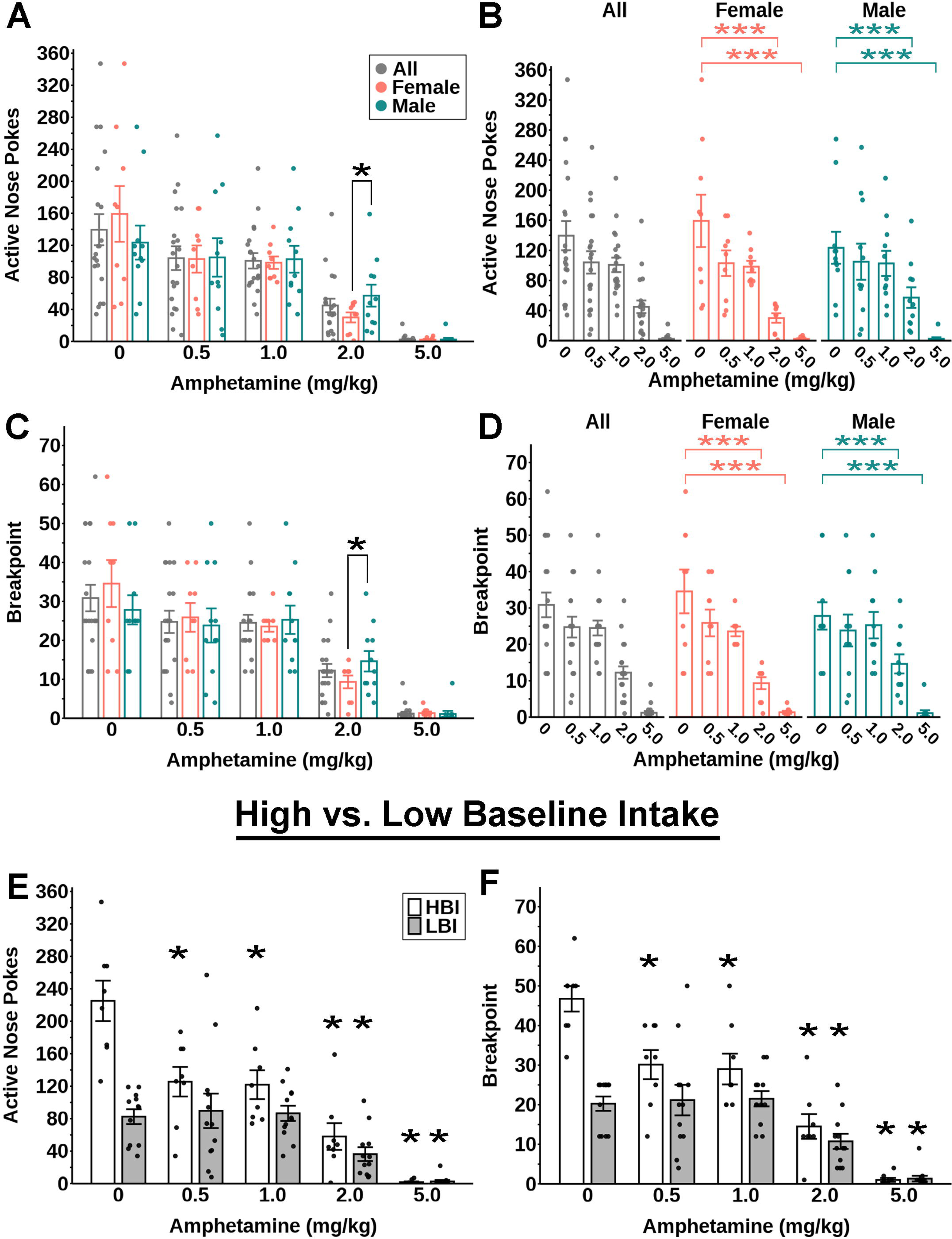
Amphetamine significantly reduced sucrose self-administration on a PR schedule. (**A-D**) Active nose pokes (**A-B**) and breakpoints (**C-D**). (**E-F**) Active nose pokes (**E**) and breakpoints (**F**) for mice in the high and low baseline sucrose intake groups. n = 20; males: n = 11; females: n = 9. High Baseline Intake group: n = 8 (n = 3 males, n = 5 females; Low Baseline Intake group: n = 12 (n = 8 males, n = 4 females). * P<0.05 vs saline; ** P<0.01 vs saline; *** P<0.001 vs saline.

We also tested whether nicotine similarly affected breakpoints or active lever presses during PR testing. As with FR3 responding, higher doses of nicotine dose-dependently decreased both break point and active lever presses (Figure 5). There was a significant effect of dose for both active nose pokes (F(5,80)=8.366, p<0.001) and breakpoint (F(5,80)=54.646, p<0.001), but no main effect of sex and no dose x sex interaction for either measure. *Post hoc* analysis showed that the highest doses of nicotine (2 mg/kg and 5 mg/kg) significantly reduced both active nose pokes and breakpoints compared to control (Figure 5A, Figure 5C). While nicotine had an effect on inactive nose pokes in the PR tests (F(5,80)=5.698, p<0.001), inactive nose pokes averaged 2.2 across all trials, once again suggesting that this is not physiologically significant. There was also no sex differences observed for any measure. When we separated the data by sex, there was a significant effect of dose for nose pokes and breakpoint in both males (nose pokes: (F(5,45)=5.434, p<0.001); breakpoint: (F(5,45)=6.270, p<0.001)) and females (nose pokes: (F(5,35)=3.763, p<0.01); breakpoint: (F(5,35)=4.500, p<0.01)). *Post hoc* analysis revealed that none of the doses significantly reduced nose pokes or breakpoint for males or females, however (Figure 4B, Figure 4D).

**Figure 5.**
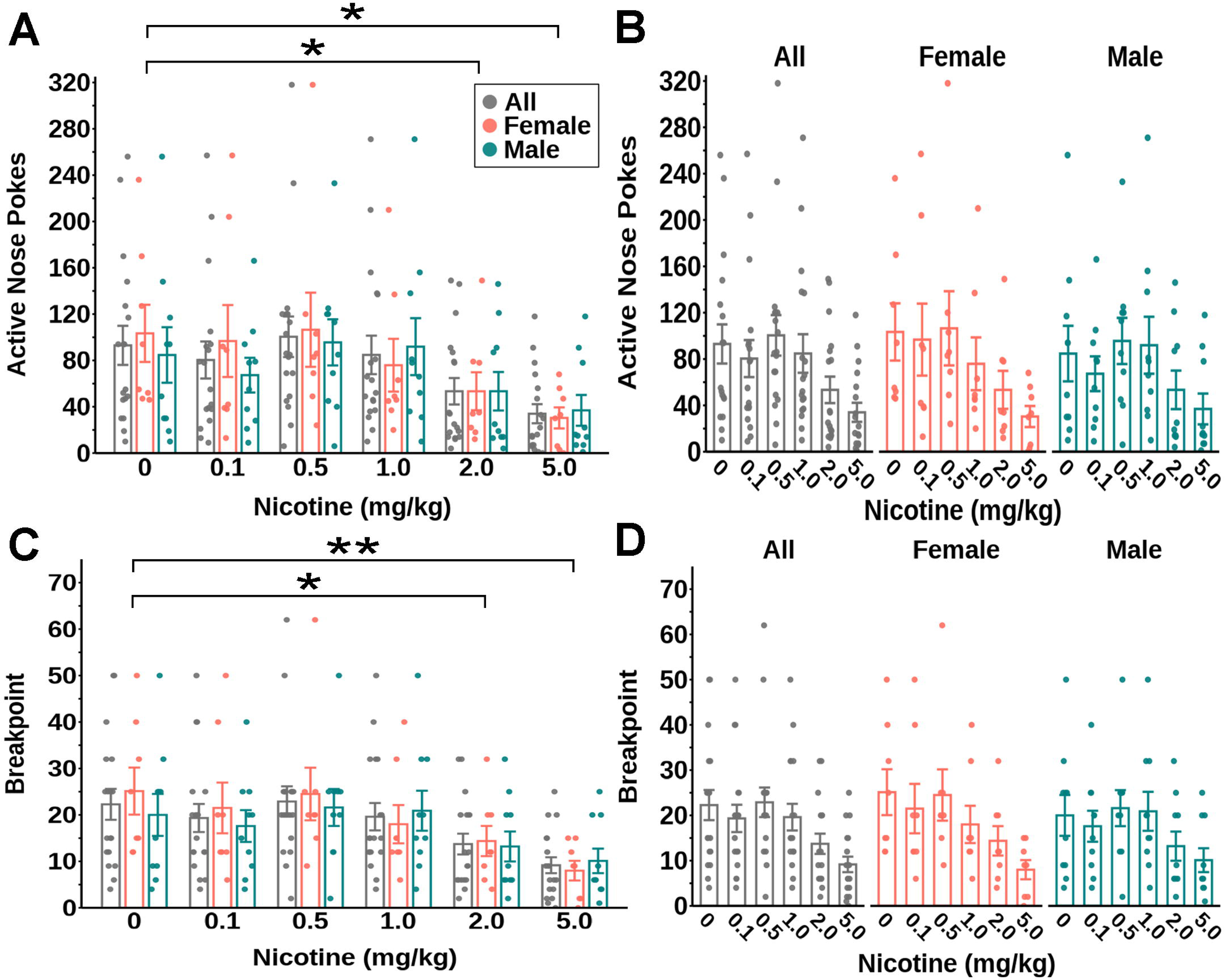
Nicotine significantly reduced sucrose self-administration on a PR schedule. Active nose pokes (**A-B**) and breakpoints (**C-D**) following nicotine treatment. All: n = 18; males: n = 10; females: n = 8. * P<0.05 vs saline; ** P<0.01 vs saline; *** P<0.001 vs saline.

### Locomotor Measurements After Nicotine Injections

We visually observed an apparent decrease in activity following injections of the highest doses of nicotine during the self-administration trials, suggesting the possibility that nicotine could have globally decreased activity to cause the observed reductions in FR3 and PR responses. To test this possibility, we next measured locomotor activity in a new cohort of mice in an open field test following treatment with the same doses of nicotine. The highest doses of nicotine did indeed decrease overall locomotor activity in these tests, as there were significant main effects of dose (F(5,50)= 21.337, p<0.001), sex (F(1,10)=6.068, p<0.05), and time (F(11,110)=5.674, p<0.001), and significant dose x time (F(55,550)=2.555, p<0.001) and sex x time Interactions (F(11,110)=1.911, p<0.05). *Post hoc* analysis showed that 5 mg/kg nicotine significantly reduced distance traveled compared to saline control, especially at early time points, and that 2 mg/kg nicotine significantly decreased distance traveled, but only at the 5 and 10 minutes (Figure 6C). Conversely, the 0.1 mg/kg dose increased distance traveled, but this was only significant for the 45-minute measurement (Figure 6C). When we separated the data by sex, for males there was a significant main effects of dose (F(5,25)=8.873, p<0.001), and time (F(11,55)=5.877, p<0.001), and significant dose x time interaction (F(55,275)=1.414, p<0.05). For females, there was a significant main effects of dose (F(5,25)=14.713, p<0.001) and significant dose x time interaction (F(55,275)=2.046, p<0.001). *Post hoc* analysis of the dose effect showed that in males only the 5 mg/kg dose reduced distance traveled compared to saline control, whereas in females the 0.5 and 5 mg/kg dose reduced distance traveled. *Post hoc* analysis of the dose x time effect showed that in males and females only the 5 mg/kg nicotine significantly reduced distance traveled compared for the first 15 minutes of the experiment when compared to saline control (Figure 6E, Figure 6F).

**Figure 6.**
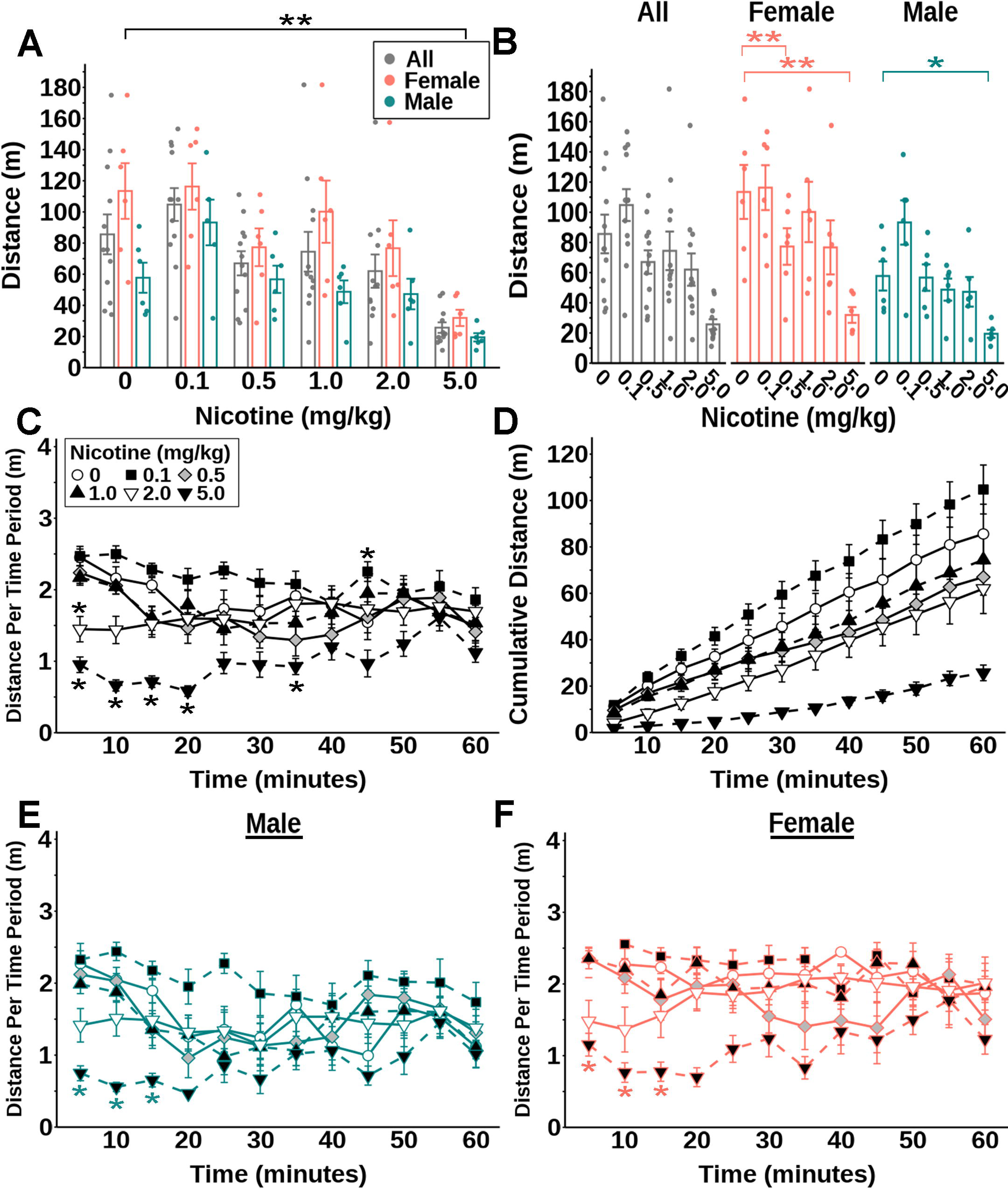
Nicotine significantly reduced locomotive behavior in an open field test. Total distance traveled (**A-B**), distance traveled per 5-minute interval (**C**), cumulative distance traveled over time (**D**), and total distance traveled per 5-minute interval for each individual male and female mouse (**E-F**) after nicotine. All: N = 12; males: n = 6; females: n = 6. * P<0.05 vs saline; ** P<0.01 vs saline; *** P<0.001 vs saline. Teal asterisks in panel E reflect significant differences vs saline for males, and orange asterisks in panel F reflect significant differences vs saline for females.

### Sucrose Free Feeding and Self-Administration in a Home Cage Setting

We had previously observed a biphasic effect of amphetamine on sucrose and palatable food intake (West et al., 2019) whereby low doses caused a delayed increase in intake compared to the rapid decrease in intake caused by higher doses. As the FR3 self-administration experiments were only 1 hour, we next tested whether nicotine or amphetamine affected longer-term sucrose intake in a home cage setting. For this experiment mice were given 24-hour access to the same sucrose pellets used in the self-administration experiments on an FR3 schedule via FED3 devices placed into their home cage. GAM analysis revealed that for home cage self-administration of sucrose on a FR3 schedule following amphetamine injections there was a 2-way interaction between time and dose, but no time x dose x baseline intake, time x dose x sex, or dose x sex interactions. *Post hoc* analysis revealed that all doses reduced sucrose self-administration, however there was a difference in length of reduction. Analysis of the effect of each dose 4 hours post injection revealed that the 2 and 5 mg/kg doses significantly reduced sucrose self-administration. Analysis of the dose x time effect over those 4 hours showed that the 0.5 and 1 mg/kg reduced self-administration for up to 1.5 hours, the 2 mg/kg doses for up to 2 hours, and the 5 mg/kg dose for up to 2.5 hours (Figure 7C). Interestingly, the 1 mg/kg dose significantly increased sucrose self-administration at the 3.5-hour mark (p < 0.05), but not at any other times (Figure 7C).

**Figure 7.**
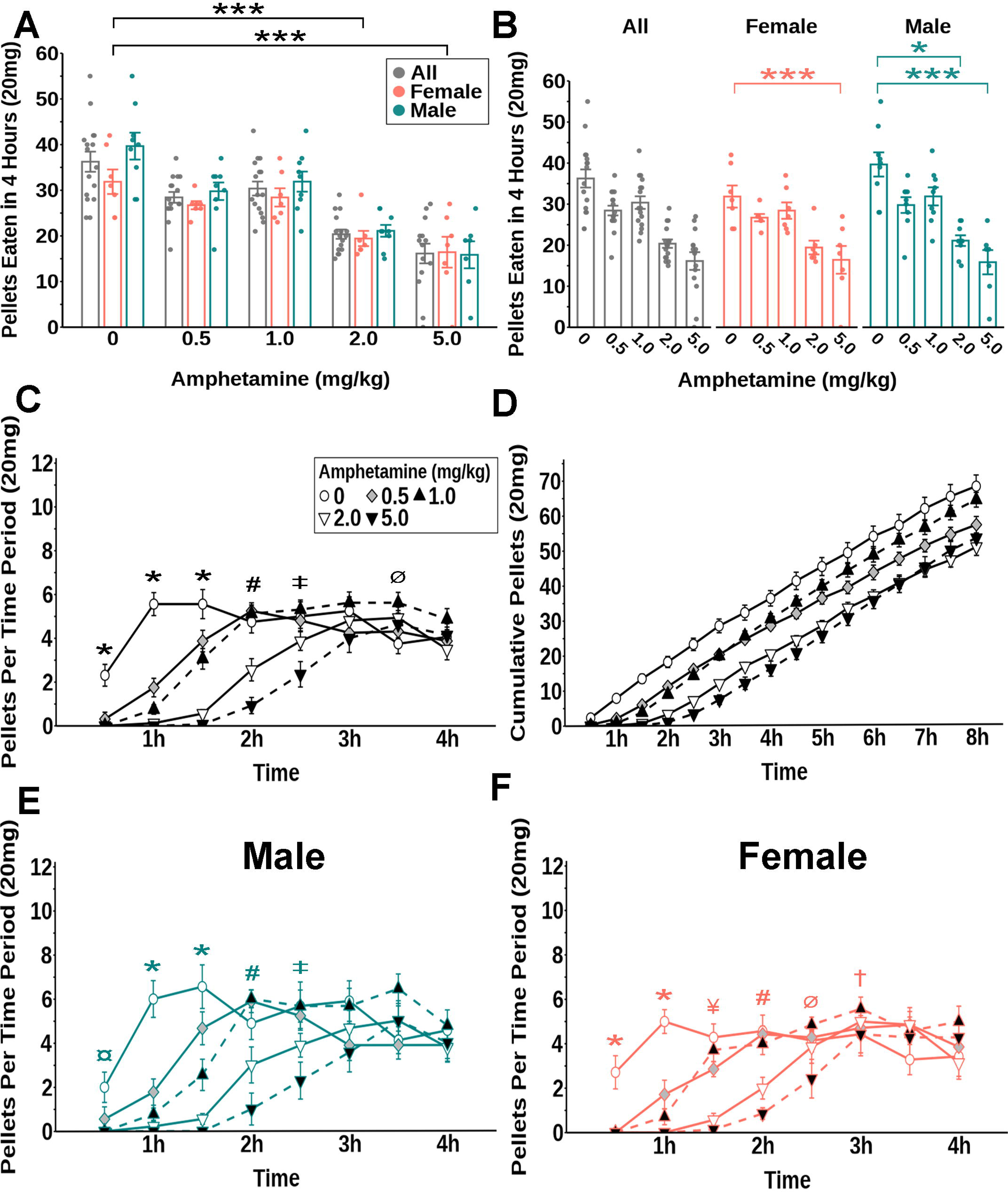
Amphetamine significantly reduced sucrose self-administration on an FR3 schedule in the home cage. (**A-B**) Total self-administered sucrose pellets eaten. (**C**) Self-administered sucrose pellets at each 30-minute interval for all mice. (**D)** Cumulative self-administered sucrose pellets for all mice over time. (**E-F**) Self-administered sucrose pellets at each 30-minute interval for male (**E**) and female (**F**) mice. n = 16; males n = 9; females n = 7. * P<0.05 vs saline; *** P<0.001 vs saline in panels A and B. * P< 0.05 vs saline (0.5, 1, 2, and 5 mg/kg); ¤ P< 0.05 vs saline (1, 2, and 5); ¥ P< 0.05 vs saline (0.5, 2, and 5); † P< 0.05 vs saline (0.5 and 1); # P< 0.05 vs saline (2 and 5); Ø P< 0.05 vs saline (1); ‡ P< 0.05 vs saline (5) in panels C, E and F. Teal symbols in panels B and E reflect significant differences vs saline for males, and orange symbols in panels B and F reflect significant differences vs saline for females.

When we separated the data by sex, *post hoc* analysis of the effect of each dose 4 hours post injection revealed that for males the 2 and 5 mg/kg doses significantly reduced sucrose self-administration, however only the 5 mg/kg doses reduced self-administration for females. Analysis of the dose x time effect over those 4 hours showed that revealed that for males, the 0.5 mg/kg dose reduced self-administration at the 1 and 1.5 hour mark, with a near significant effect at 30 minutes (p=0.0527), the 1 mg/kg reduced self-administration for up to 1.5 hours, the 2 mg/kg doses for up to 2 hours, and the 5 mg/kg dose for up to 2.5 hours (Figure 7E). No increase in self-administration was observed for males. For females, the 0.5 mg/kg dose reduced self-administration up to 1.5 hours, the 1 mg/kg reduced self-administration for up to 1 hour, the 2 mg/kg doses for up to 2 hours, and the 5 mg/kg dose for up to 2 hours with a near significant decrease at 2.5 hours (p=0.0577). The 0.5 mg/kg dose increased self-administration at the 3-hour mark and the 1 mg/kg dose also increased self-administration at 2.5 and 3 hours (Figure 7F).

We also tested whether amphetamine affected free intake of sucrose in the home cage. For these experiments, sucrose pellets were delivered *ad libitum* via FED3 devices in their home cages (i.e. no operant requirement). GAM analysis revealed that there was a 2-way interaction between time and dose, but no time x dose x baseline intake, time x dose x sex, or dose x sex interactions. *Post hoc* analysis revealed that all doses reduced sucrose self-administration, but there was a difference in length of reduction. Analysis of the effect of each dose 4 hours post injection revealed that only the 5 mg/kg dose significantly reduced sucrose self-administration. Analysis of the dose x time effect over those 4 hours showed that the 0.5 and 1 mg/kg decreased sucrose intake for up to 1 hour. Whereas the 2 mg/kg dose reduced sucrose intake for up to 1.5 hours, and the 5 mg/kg dose for up to 2.5 hours (Figure 8C). Unlike in the self-administration condition, there was no significant increase in sucrose intake at any time (Figure 8C). When we separated the data by sex, *post hoc* analysis of the effect of each dose 4 hours post injection revealed that for males and females the 5 mg/kg dose significantly reduced sucrose self-administration. Analysis of the dose x time effect over those 4 hours showed that for males, the 0.5 and 1 mg/kg doses reduced sucrose intake for up to 1 hour and the 2 and 5 mg/kg doses for up to 1.5 hours. The 5 mg/kg dose also decreased intake at 3 hours (Figure 8E). For females, the 1 and 2 mg/kg reduced self-administration for up to 1 hour and the 5 mg/kg dose for up to 2.5 hours (Figure 8F).

**Figure 8.**
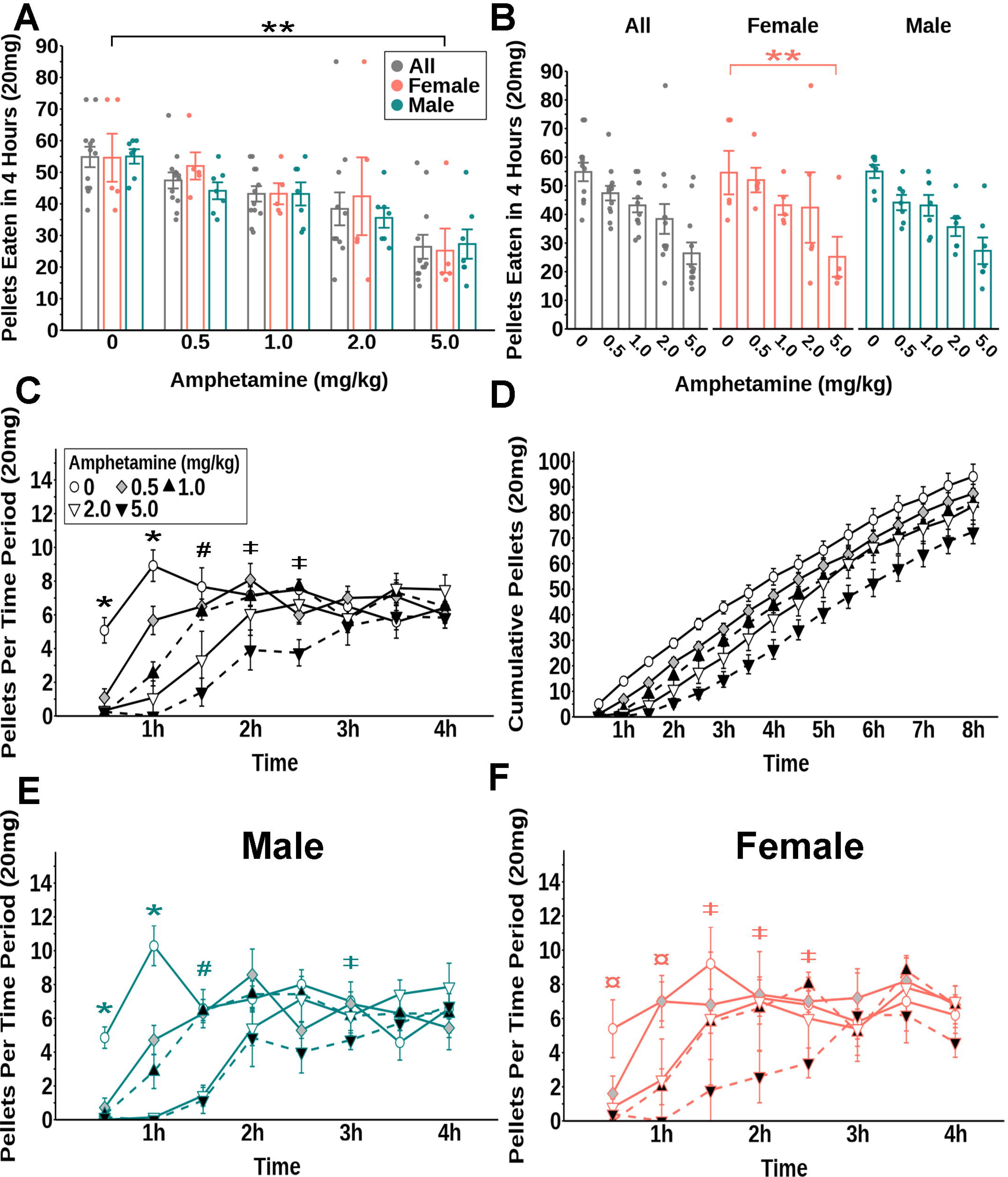
Amphetamine significantly reduced free access sucrose intake in the home cage. (**A-B**) Total sucrose pellets obtained. (**C**) Sucrose pellets obtained for all mice at each 30-minute interval. (**D**) Cumulative sucrose pellets obtained for all mice over time. (**E-F**) Sucrose pellets obtained at each 30-minute interval for males and females. n = 12; males n = 7; females n = 5. ** P<0.01 vs saline in panels A and B. * P< 0.05 vs saline (0.5, 1, 2, and 5 mg/kg); ¤ P< 0.05 vs saline (1, 2, and 5); # P< 0.05 vs saline (2 and 5); ‡ P< 0.05 vs saline (5) in panels C, E and F. Teal symbols in panel B and E reflect significant differences vs saline for males, and orange symbols in panel B and F reflect significant differences vs saline for females.

GAM analysis revealed that for nicotine’s effects on sucrose self-administration there was a 2-way interaction between time and dose, but no time x dose x baseline intake, time x dose x sex, or dose x sex interactions. *Post hoc* analysis of the effect of each dose 2 hours post injection revealed that only the 5 mg/kg dose significantly reduced sucrose self-administration. Analysis of the dose x time effect over those 2 hours showed that the 5 mg/kg dose decreased sucrose self-administration for up to an hour, the 2 mg/kg had a near significant decrease (p=0.05) at 1 hour, and no significant reduction in sucrose self-administration was seen in any of the other doses (Figure 9C). When we separated the data by sex, *post hoc* analysis of the effect of each dose 2 hours post injection revealed that the 5 mg/kg dose significantly reduced sucrose self-administration for females only. Analysis of the dose x time effect over those 2 hours showed that the 5 mg/kg dose reduced self-administration for up to 1 hour in both males and females and the 2 mg/kg dose reduced self-administration at the 1-hour mark in males only (Figure 9E), and the 0.5 mg/kg dose increased self-administration at 30 minutes in females only (Figure 9F).

**Figure 9.**
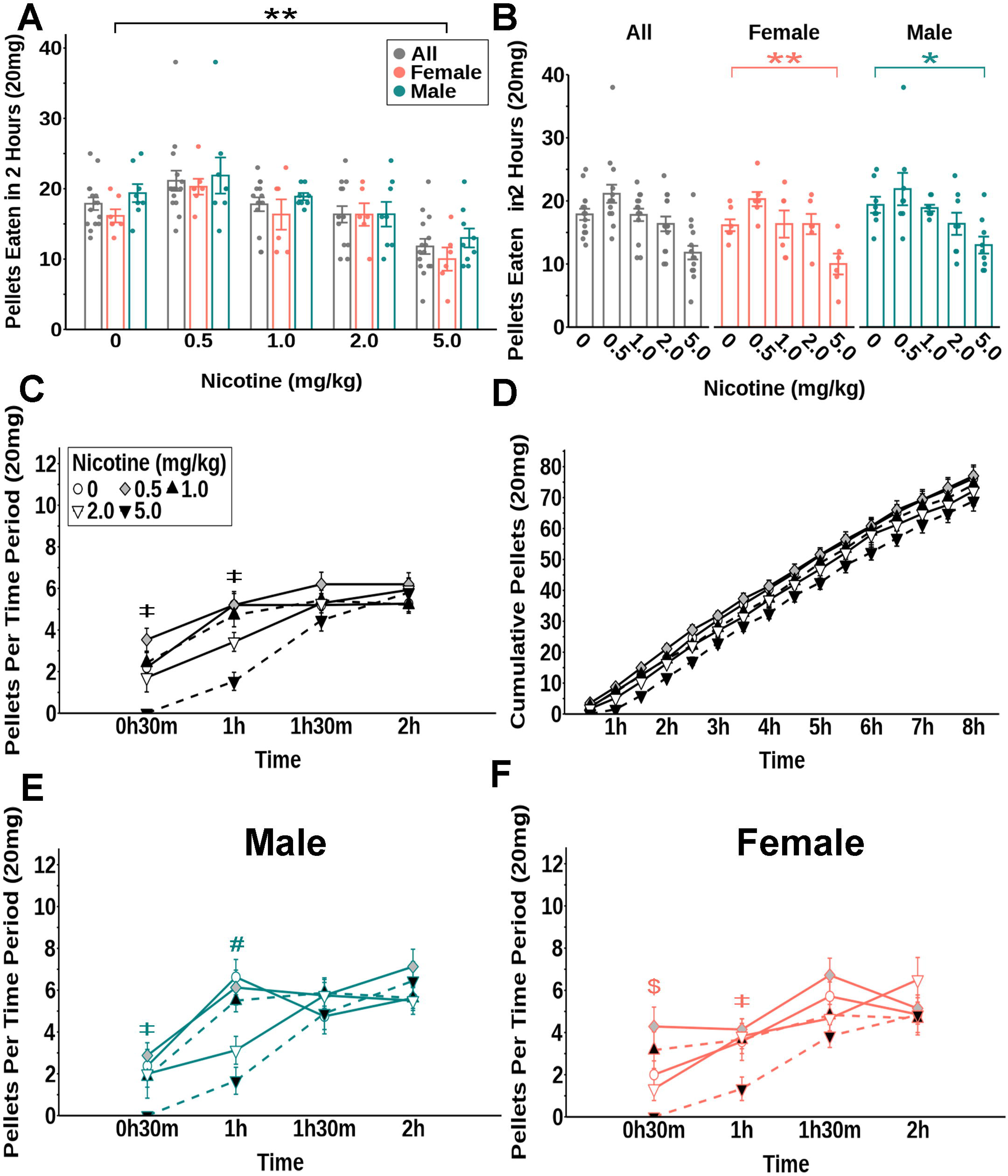
Nicotine significantly reduced sucrose self-administration on an FR3 schedule in the home cage. (**A-B**) Total sucrose pellets obtained. (**C**) Sucrose pellets obtained at each 30-minute interval for all mice. (**D**) Cumulative sucrose pellets obtained over time for all mice. (**E-F**) Sucrose pellets obtained at each 30-minute interval for male (**E**) and female (**F**) mice. All: n = 17; males n = 9; females n = 8. * P<0.05 vs saline; ** P<0.01 vs saline in panels A and B. $ P< 0.05 vs saline (0.5 and 5 mg/kg); # P< 0.05 vs saline (2 and 5); ‡ P< 0.05 vs saline (5) in panels C, E and F. Teal symbols in panel B and E reflect significant differences vs saline for males, and orange symbols in panel B and F reflect significant differences vs saline for females.

For nicotine’s effects on free feeding sucrose, GAM analysis revealed that there were time x dose and sex x dose interactions, but no time x dose x baseline intake or time x dose x sex interactions. *Post hoc* analysis of the dose x time interaction revealed that the 1 and 5 mg/kg dose decreased sucrose intake for 30 minutes. No significant reduction in sucrose intake was seen in any of the other doses, however, the 2 mg/kg dose did significantly increase sucrose intake from 1.5 to 2 hours post nicotine injection (Figure 10C). When we separated the data by sex, *Post hoc* analysis revealed that for males, the 5 mg/kg dose reduced intake for 30 minutes and the 2 mg/kg dose increased self-administration at the 2-hour mark (Figure 10E). For females, none of the doses has any significant effect on sucrose intake in the 2-hour window analyzed (Figure 10F).

**Figure 10.**
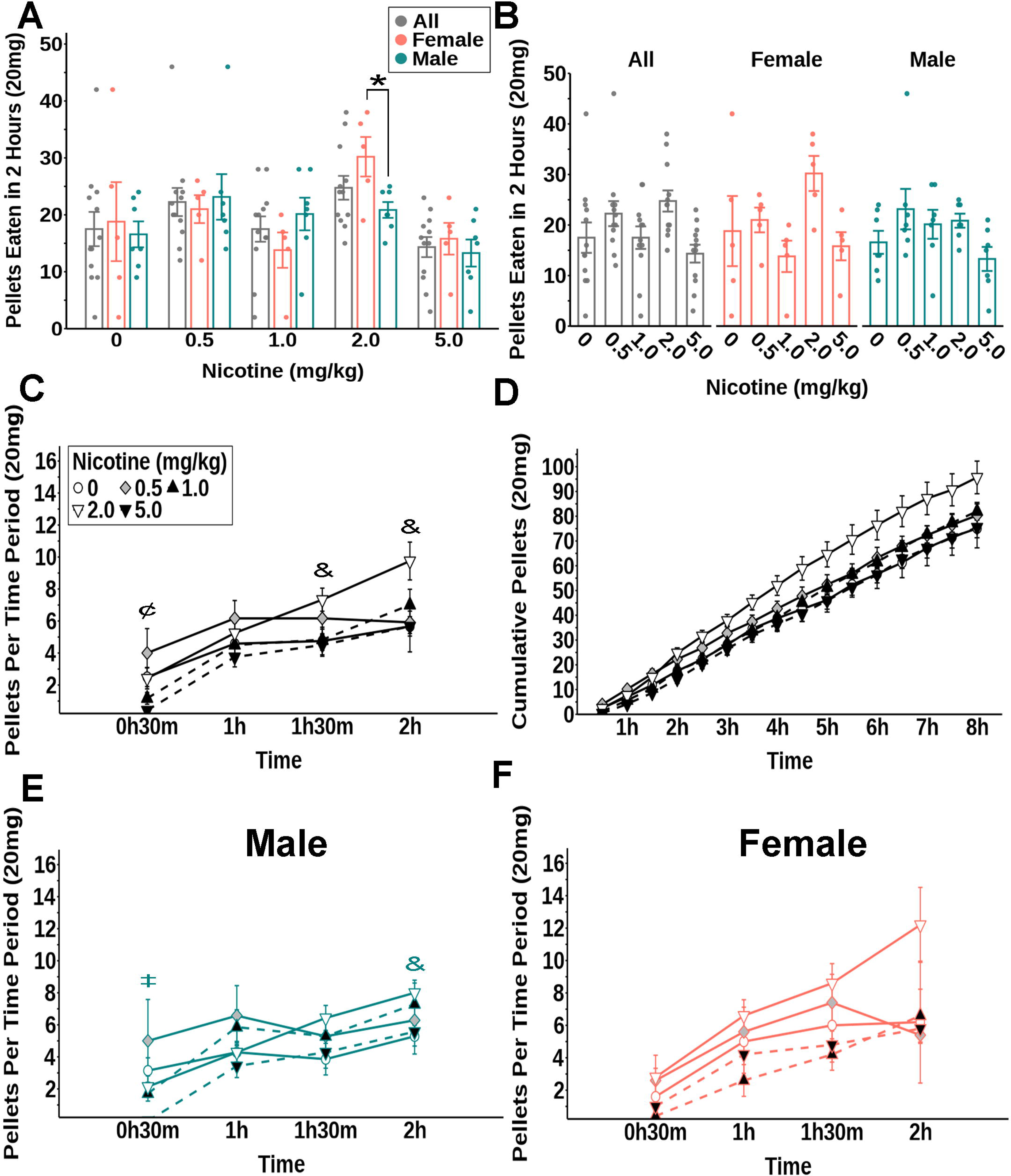
Nicotine significantly altered free access sucrose intake in the home cage. (**A-B**) Sucrose pellets obtained. (**C**) Sucrose pellets obtained for all mice at each 30-minute interval. (**D**) Cumulative sucrose pellets obtained for all mice over time. (**E-F**) Sucrose pellets obtained for males and females at each 30-minute interval. All: n = 12; males n = 7; females n = 5. * P<0.05 vs saline in Panel A. ¢ P< 0.05 vs saline (0.5 and 5 mg/kg); & P< 0.05 vs saline (2); ‡ P< 0.05 vs saline (5) in panels C and E. Teal symbols in panel E reflect significant differences vs saline for males, and orange symbols in panel F reflect significant differences vs saline for females.

## Discussion

In these studies, we tested whether amphetamine and nicotine decreased sucrose intake in both operant chamber self-administration, extended access home-cage self-administration, and free intake models. Amphetamine showed a strong dose-dependent effect in its ability to reduce sucrose self-administration in both operant chambers and the home cage as well as free intake. In contrast, only the highest dose of nicotine decreased sucrose self-administration and free intake but there did seem to be some variability in the effects of lower doses of nicotine on free intake. The effects of nicotine may have been due, at least in part, to a reduction in general locomotor behaviors, as the doses that decreased sucrose self-administration also significantly inhibited general locomotor activity, although there were some differences in the time courses of these two effects. Finally, overall, there were no sex differences in the responses to amphetamine or nicotine in these studies.

The results presented here indicate that amphetamine inhibited feeding to a similar degree when tested under normal, baseline, *ad libitum* free feeding conditions and during operant conditioning self-administration assays, with some minor variances in the duration of their effects (Figure 7, Figure 8). This primarily holds true for nicotine as well, although there was some indication that *ad libitum* free feeding may have been more sensitive to lower doses of nicotine (Figure 9, Figure 10). This suggests that amphetamine’s effect on normal baseline feeding and the motivational and rewarding aspects of feeding may be mediated by similar mechanisms. These results are also consistent with other studies in the literature showing that amphetamine is able to effectively decrease feeding under normal ad libitum feeding conditions (Cannon et al., 2004; Sills & Vaccarino, 1996; Winn et al., 1982; Dobrzanski & Doggett, 1976).

One interesting observation from these studies is the variability in the response to low doses of amphetamine. Although low doses of amphetamine significantly inhibited sucrose intake under both free feeding and FR3 conditions in an extended access home cage setting, these same low doses appeared to be ineffective when tested in operant chambers. Further analysis of the operant chamber revealed that the ability of amphetamine to decrease active nose pokes and pellets eaten depended on the baseline level of responding of the mice, with mice showing high levels of baseline active nose pokes sensitive to amphetamine but those with low baseline activity were insensitive. One possible explanation for these differential responses is that the differences in sucrose access between the two conditions contributed to the differences in intake. Mice were only given intermittent access to sucrose during the 1-hour trial each day for the operant chamber experiments, whereas the home-cage mice had continuous, *ad libitum*, 24 hour access to the sucrose pellets. The intermittent access to a palatable substance in the operant chambers may have induced binge-like behaviors where mice will consume a higher portion of their daily calories during this limited sucrose access window and show increased reinforcement for palatable foods (Berner et al., 2008; Avena et al., 2008; Wojnicki et al., 2010). This is in contrast to the continuous sucrose access in the home cage which has been shown to decrease motivation for sucrose reward (Finnell & Ferrario, 2022). Another possibility is that these differences resulted from differences in dopamine dynamics in the nucleus accumbens (NAc) following amphetamine administration. Although amphetamine causes the release of several different monoamines including dopamine, norepinephrine, and serotonin, its anorexigenic effects are believed to be mediated by dopamine release in the NAc (Cannon et al., 2004; Chen et al., 2001). Microdialysis studies have shown that NAc dopamine levels following amphetamine administration vary depending on the baseline sucrose intake where rats with low baseline intake show diminished NAc dopamine in response to AMPH compared to rats with high baseline intake (Sills & Crawley, 1996). Thus, it is possible that the differences in the responses to low dose amphetamine were due to differences in NAc dopamine release between the two groups.

In addition to the variability in the effects of low dose amphetamine, it was somewhat surprising that there were no increases in sucrose intake after low dose amphetamine in any of these experiments. We previously found that low dose amphetamine (0.5 mg/kg) increased sucrose intake in a two-bottle choice test and tended to increase intake of a palatable high fat/high sugar food in a binge intake model (West et al., 2019). Furthermore, other studies have shown that low doses of amphetamine can also increase food intake, however there is some variability in which doses cause this effect. For example, some studies have shown that 0.5 mg/kg decreased or did not affect feeding while doses lower than 0.5 mg/kg increased food intake (Sills & Vaccarino, 1996; Winn et al., 1982; Evans & Vaccarino, 1990; Orthen-Gambill, 1985). It has also been shown that the effects of low doses of amphetamine may depend on the satiety level of the subject. One study found that amphetamine sulphate will increase feeding during the light phase when mice are typically sated and decreases feeding during the dark phase when mice consume most of their daily calories (Dobrzanski & Doggett, 1976). Therefore, it is possible that the dose by which amphetamine is able to increase feeding may depend on a number of conditions including experimental methodologies, the type of food reward, the time of treatment, and the physiological state of the subject, which could help to explain the lack of an increase in feeding observed in these studies.

Nicotine was also able to reduce sucrose-self administration in these experiments, but only at high doses of nicotine, 2 and 5 mg/kg, (Figure 3, Figure 5, Figure 9, Figure 10) that have been reported to have motor and other side effects. Our results were confounded by the finding that the same doses of nicotine also significantly depressed locomotion in open field tests. Thus, it is possible that these alterations in locomotor activity could have impeded mice’s ability to nose poke for sucrose (Figure 6). While the highest doses of nicotine used reduce locomotive behaviors on a similar timescale as the reduction seen in sucrose self-administration (Figure 3C, Figure 6C), the duration of inhibition of locomotion and sucrose self-administration after high dose nicotine differed across the two experiments. The depression of locomotor activity lasted longer than the reduction in food intake in the FR3 operant conditioning trials (Figure 3D, Figure 6D), suggesting the possibility that all the effects of sucrose intake in the self-administration trials may not be fully accounted for by changes in locomotion.

These studies also appear to contradict some other studies showing that acute nicotine increased operant responding and breakpoint for food reward or food associated cues (Palmatier et al., 2009; Popke et al., 2000; Brunzell et al., 2006). It should be noted, however, that in all studies showing increases in operant responding, mice were food restricted for the entirety of the experiment and Donney et al. have reported that the enhanced food reinforcement effect of nicotine is primarily observed under conditions of food restriction and not under conditions of *ad libitum* feeding (Donny et al., 2011). Thus, the differences observed here may be due to the nature of the feeding paradigms employed in these studies. This is further demonstrated by our results showing that specific doses of nicotine may increase food intake depending on the type of food access (self-administration vs. free access) (Figure 9C, Figure 10C).

In contrast to robust sex differences observed in many other drug responses as well as the effects of manipulation of multiple systems controlling feeding and body weight, we only observed minor sex differences in the effects of either amphetamine or nicotine in these experiments. We observed that lower doses of amphetamine may have a greater effect on sucrose motivation in females than in males although the same doses of amphetamine decreased this motivation for both males and females. This suggests that there may be sex differences in the sensitivity of different amphetamine doses but not in their qualitative effects. This finding is somewhat surprising in light of the robust sex differences in the control of food intake and body weight (Morio et al., 1997; Carpenter et al., 1998; Romanski et al., 2000; Hellström et al., 1996; Leibel & Hirsch, 1987; Geary et al., 1994; Clegg et al., 2007; Rosenbaum et al., 1996; Valenstein et al., 1967; Piggott, 1979; Klump et al., 2013; Munro et al., 2006; Sinclair et al., 2017) and the rewarding, reinforcing, motivational, and locomotor sensitizing qualities of both of these drugs (Zakiniaeiz et al., 2019; Milesi-Hallé et al., 2007; Kawa & Robinson, 2019; Donny et al., 2000; Wang et al., 2014; Locklear et al., 2012; Lynch, 2009; Leventhal et al., 2007). This suggests that although some of the mechanisms responsible for the rewarding and reinforcing effects of amphetamine and nicotine may overlap with those for their control of feeding, the primary mechanisms mediating these effects are likely distinct. This possibility is supported by other studies which demonstrate both distinct and overlapping brain mechanisms mediating drug reinforcement and food reward. For example, dopamine depletion in the nucleus accumbens, which impairs cocaine self-administration, has little effect on food reinforced responding (Salamone et al., 2003). Furthermore, D1R and D2R antagonists administered to the nucleus accumbens shell decrease cocaine but not food reinforcement (Di Ciano et al., 2003; Bari & Pierce, 2005). However, these same antagonists applied to the nucleus accumbens core decrease responding for both food and drugs (Bari & Pierce, 2005). Finally, GABA-B receptor agonists at low concentrations decrease nicotine self-administration, but not food responding whereas high concentrations of the agonist decrease both (Paterson et al., 2005). Thus, it appears likely that the mechanisms mediating amphetamine and nicotine anorexia may contain both distinct and overlapping mechanisms from those mediating their rewarding, reinforcing, and motivational effects, and further identifying how these effects are mediated may allow for the development of new approaches to decrease feeding. Even though we did not detect many interactions between dose and sex in the experiments shown here, when we separated the data by sex and ran the analysis, we noted subtle differences in which doses were effective if they were effective at all. However, as these effects were not consistent across experiments, it is unclear what this means.

Although amphetamine and nicotine had clear effects on feeding under the multiple different conditions studied here, there were some limitations that should be considered when interpreting these results. First, mice were single housed for the duration of all experiments. This allowed for measurement of food intake of individual mice and was essential for the acquisition of operant conditioning sucrose self-administration. Others have shown that single housing can impact reward responding and behavioral responses to amphetamine and nicotine, however (Jones et al., 1990; Cheeta et al., 2001). All experiments in this study were also performed at the beginning of the dark phase to increase acquisition and optimize the performance of the mice in the operant conditioning paradigm, which is in contrast to many other studies testing the effects of nicotine and amphetamine. Thus, this raises potential issues when comparing these results to other studies. One final potential limitation is that this study focused exclusively on the acute effects of nicotine. Most other studies of nicotine’s effects on feeding and body weight have focused on chronic administration, which more similarly mimics the human condition. These studies have provided important information on the acute effects of nicotine that can serve as a foundation for other future experiments testing the effects of chronic administration.

In summary, we have demonstrated here that amphetamine and nicotine reduced both sucrose self-administration and the acute intake of sucrose during *ad libitum* free-feeding access. Unlike the broader and more robust, rewarding, reinforcing, and locomotor effects of amphetamine and nicotine, there were only minor sex differences in their effects of on sucrose self-administration or free intake. Finally, we demonstrated that the effects of low dose amphetamine (0.5 mg/kg) depended on the baseline intake of sucrose and did not cause the delayed increase in intake that we had observed previously.

