## Supplemental Information for "Amphetamine and Nicotine Reduce Sucrose Self-Administration Independent of Sex"

**Model Selection**

**GAM of nose pokes in PR trials following amphetamine treatment**

When designing our model, we chose to use the maximum likelihood estimation method so that comparison of several models for best fit could be performed. We also chose to allow for automatic selection of smooth terms by setting select = TRUE in the mgcv package. We first started by testing for overdispersion in our model by using the dispersion test from the DARHMa package and found significant overdispersion of our data when using a Poisson distribution family (p<0.01). Therefore, the negative binomial family was tested to account for overdispersion. Furthermore, to determine which variables significantly accounted for the variance found in our data we looked at five models under the negative binomial family. The models were-

Model 1: Nose Pokes ~ Dose + s(Mouse, bs = "re")

Model 2: Nose Pokes ~ Dose * Sex + s(Mouse, bs = "re")

Model 3: Nose Pokes ~ Dose * Baseline Intake + s(Mouse, bs = "re")

Model 4: NosePokes ~ Dose * IntakeRating + Dose * Sex + s(Mouse, bs = "re")

Model 5: Nose Pokes ~ s(Time, by = Dose), k = 8) + Dose * Sex + s(Mouse, bs = "re").

Likelihood ratio testing for variable selection primarily used ANVOA tests. This analysis showed that Model 4 performed significantly better than Model 1, Model 2, and Model 3 (p<0.05). Therefore, Model 4 was selected. Model 1 accounted for 84.3% of the deviance in the data.

**GAM of breakpoint in PR trials following amphetamine treatment**

When designing our model, we chose to use the maximum likelihood estimation method so that comparison of several models for best fit could be performed. We also chose to allow for automatic selection of smooth terms by setting select = TRUE in the mgcv package. We first started by testing for overdispersion in our model by using the dispersion test from the DARHMa package and found significant overdispersion of our data when using a Poisson distribution family (p<0.01). Therefore, the negative binomial family was tested to account for overdispersion. Furthermore, to determine which variables significantly accounted for the variance found in our data we looked at five models under the negative binomial family. The models were-

Model 1: Breakpoint ~ Dose + s(Mouse, bs = "re")

Model 2: Breakpoint ~ Dose * Sex + s(Mouse, bs = "re")

Model 3: Breakpoint ~ Dose * Baseline Intake + s(Mouse, bs = "re")

Model 4: Breakpoint ~ Dose * IntakeRating + Dose * Sex + s(Mouse, bs = "re")

Model 5: Nose Pokes ~ s(Time, by = Dose), k = 8) + Dose * Sex + s(Mouse, bs = "re").

Likelihood ratio testing for variable selection primarily used ANVOA tests. This analysis showed that Model 4 performed significantly better than Model 1, Model 2, and Model 3 (p<0.05). Therefore, Model 4 was selected. Model 1 accounted for 87.5% of the deviance in the data.

**GAM of self-administered pellets in home cage FR3 self-administration trials following amphetamine treatment**

When designing our model, we chose to use the maximum likelihood estimation method so that comparison of several models for best fit could be performed. We also chose to allow for automatic selection of smooth terms by setting select = TRUE in the mgcv package. We first started by testing for overdispersion in our model by using the dispersion test from the DARHMa package and found significant overdispersion of our data when using a Poisson distribution family (p<0.01). Therefore, the negative binomial family was tested to account for overdispersion. Furthermore, to determine which variables significantly accounted for the variance found in our data we looked at four models under the negative binomial family. The models were-

Model 1: Nose Pokes ~ s(Time, by = Dose, k = 8) + Dose + Sex + s(Mouse, bs = "re")

Model 2: Nose Pokes ~ s(Time, by = interaction(Dose, Baseline Intake), k = 8) + Baseline Intake * Dose + Sex + s(Mouse, bs = "re")

Model 3: Nose Pokes ~ s(Time, by = interaction(Dose, Sex), k=8) + Dose * Sex + s(Mouse, bs = "re").

Model 4: Nose Pokes ~ s(Time, by = Dose), k = 8) + Dose * Sex + s(Mouse, bs = "re").

Likelihood ratio testing for variable selection primarily used ANVOA tests. This analysis showed that Model 1 performed comparatively to more complex models with no significant differences observed when compared to Model 3 (p = 1.00) and Model 4 (p = 0.669). When examining Model 2, BIC analysis was used due to missing datapoints which did not allow for ANOVA likelihood ratio testing. This test indicated that Model 1 performed better than Model 2. As this was the case, and Model 1 is the simpler model, Model 1 was selected. Model 1 accounted for 63.2% of the deviance in the data. Due to heteroscedasticity detected in the residuals of this model, Huber-White standard errors were used for robustness.

**GAM of free access pellets received in home cage trials following amphetamine treatment**

When designing our model, we chose to use the maximum likelihood estimation method so that comparison of several models for best fit could be performed. We also chose to allow for automatic selection of smooth terms by setting select = TRUE in the mgcv package. We first started by testing for overdispersion in our model by using the dispersion test from the DARHMa package and found significant overdispersion of our data when using a Poisson distribution family (p<0.01). Therefore, the negative binomial family was tested to account for overdispersion. Furthermore, to determine which variables significantly accounted for the variance found in our data we looked at four models under the negative binomial family. The models were-

Model 1: Nose Pokes ~ s(Time, by = Dose, k = 8) + Dose + Sex + s(Mouse, bs = "re")

Model 2: Nose Pokes ~ s(Time, by = interaction(Dose, Baseline Intake), k = 8) + Baseline Intake * Dose + Sex + s(Mouse, bs = "re")

Model 3: Nose Pokes ~ s(Time, by = interaction(Dose, Sex), k=8) + Dose * Sex + s(Mouse, bs = "re").

Model 4: Nose Pokes ~ s(Time, by = Dose), k = 8) + Dose * Sex + s(Mouse, bs = "re").

Likelihood ratio testing for variable selection used ANVOA tests. This analysis showed that Model 1 performed comparatively to more complex models with no significant differences observed when compared to Model 2 (p = 0.168), Model 3 (p = 0.978), and Model 4 (p = 0.320). As this was the case, and Model 1 is the simpler model, Model 1 was selected. Model 1 accounted for 52.7% of the deviance in the data. Due to heteroscedasticity detected in the residuals of this model, Huber-White standard errors were used for robustness.

**GAM of free access pellets received in home cage trials following nicotine treatment**

When designing our model, we chose to use the maximum likelihood estimation method so that comparison of several models for best fit could be performed. We also chose to allow for automatic selection of smooth terms by setting select = TRUE in the mgcv package. We first started by testing for overdispersion in our model by using the dispersion test from the DARHMa package and found significant overdispersion of our data when using a Poisson distribution family (p<0.01). Therefore, the negative binomial family was tested to account for overdispersion. Furthermore, to determine which variables significantly accounted for the variance found in our data we looked at three models under the negative binomial family. The models were-

Model 1: Sucrose Pellets ~ s(Time, by = Dose, k = 8) + Dose + Sex + s(Mouse, bs = "re")

Model 2: Sucrose Pellets ~ s(Time, by = interaction(Dose, Sex), k=8) + Dose * Sex + s(Mouse, bs = "re").

Model 3: Sucrose Pellets ~ s(Time, by = Dose), k = 8) + Dose * Sex + s(Mouse, bs = "re").

Likelihood ratio testing for variable selection used ANVOA tests. This analysis showed that Model 1 performed comparatively to more complex models with no significant differences observed when compared to Model 2 (p = 0.651) and Model 3 (p = 0.765). As this was the case, and Model 1 is the simpler model, Model 1 was selected. Model 1 accounted for 52.7% of the deviance of the data. Due to heteroscedasticity detected in the residuals of this model, Huber-White standard errors were used for robustness.

**GAM of free access pellets received in home cage trials following nicotine treatment**

When designing our model, we chose to use the maximum likelihood estimation method so that comparison of several models for best fit could be performed. We also chose to allow for automatic selection of smooth terms by setting select = TRUE in the mgcv package. We first started by testing for overdispersion in our model by using the dispersion test from the DARHMa package and found significant overdispersion of our data when using a Poisson distribution family (p<0.01). Therefore, the negative binomial family was tested to account for overdispersion. Furthermore, to determine which variables significantly accounted for the variance found in our data we looked at three models under the negative binomial family. The models were-

Model 1: Nose Pokes ~ s(Time, by = Dose, k = 8) + Dose + Sex + s(Mouse, bs = "re")

Model 2: Nose Pokes ~ s(Time, by = interaction(Dose, Sex), k=8) + Dose * Sex + s(Mouse, bs = "re").

Model 3: Nose Pokes ~ s(Time, by = Dose), k = 8) + Dose * Sex + s(Mouse, bs = "re").

Likelihood ratio testing for variable selection used ANVOA tests. This analysis showed that Model 3 performed significantly better than Model 1 (p<0.05) but did not perform better than Model 2 (p = 0.123). As this was the case, and Model 3 is the simpler model, Model 3 was selected. Due to heteroscedasticity detected in the residuals of this model, Huber-White standard errors were used for robustness.

| **Supplemental Table 1** |  |  |  |  |  |  |  |
| --- | --- | --- | --- | --- | --- | --- | --- |
| **Summary of GAM Results for Active Nose pokes in Amphetamine Treatment Self-Administration Progressive Ratio Trials** | | | | | | | |
| **Effect** | **Estimate*** | **Std. Error*** | **t-value** | **Type Of Term** | **EDF** | **F-value** | **p-value** |
| (Intercept) | 5.358 | 0.247 | 21.651 | Linear |  |  | 0.000 |
| Dose0.5 | -0.589 | 0.260 | -2.269 | Linear |  |  | 0.023 |
| Dose1 | -0.624 | 0.260 | -2.405 | Linear |  |  | 0.016 |
| Dose2 | -1.732 | 0.265 | -6.531 | Linear |  |  | 0.000 |
| Dose5 | -4.713 | 0.387 | -12.168 | Linear |  |  | 0.000 |
| IntakeRatingLow | -1.089 | 0.300 | -3.635 | Linear |  |  | 0.000 |
| SexM | 0.170 | 0.296 | 0.575 | Linear |  |  | 0.565 |
| Dose0.5:IntakeRatingLow | 0.596 | 0.315 | 1.893 | Linear |  |  | 0.058 |
| Dose1:IntakeRatingLow | 0.749 | 0.314 | 2.383 | Linear |  |  | 0.017 |
| Dose2:IntakeRatingLow | 0.400 | 0.321 | 1.246 | Linear |  |  | 0.213 |
| Dose5:IntakeRatingLow | 1.378 | 0.470 | 2.933 | Linear |  |  | 0.003 |
| Dose0.5:SexM | -0.166 | 0.311 | -0.534 | Linear |  |  | 0.593 |
| Dose1:SexM | -0.108 | 0.310 | -0.348 | Linear |  |  | 0.728 |
| Dose2:SexM | 0.530 | 0.318 | 1.666 | Linear |  |  | 0.096 |
| Dose5:SexM | -0.544 | 0.455 | -1.195 | Linear |  |  | 0.232 |
| s(Mouse) |  |  |  | Random Effect | 13.152 | 68.725 | 0.000 |
| **Model Statistics** |  |  |  |  |  |  |  |
| BIC | 994.900 |  |  |  |  |  |  |
| R² (Deviance Explained) | 0.843 |  |  |  |  |  |  |
| Random Effect Variance (Mouse) | 0.422 |  |  |  |  |  |  |
| * on log scale |  |  |  |  |  |  |  |
| **Supplemental Table 2**  **Summary of GAM Results for Breakpoints in Amphetamine Treatment Self-Administration Progressive Ratio Trials** | | | | | | | |
| **Effect** | **Estimate*** | **Std. Error*** | **t-value** | **Type Of Term** | **EDF** | **F-value** | **p-value** |
| (Intercept) | 3.822 | 0.156 | 24.579 | Linear |  |  | 0.000 |
| Dose0.5 | -0.437 | 0.149 | -2.931 | Linear |  |  | 0.003 |
| Dose1 | -0.525 | 0.150 | -3.496 | Linear |  |  | 0.000 |
| Dose2 | -1.422 | 0.176 | -8.103 | Linear |  |  | 0.000 |
| Dose5 | -3.720 | 0.399 | -9.323 | Linear |  |  | 0.000 |
| IntakeRatingLow | -0.885 | 0.192 | -4.597 | Linear |  |  | 0.000 |
| SexM | 0.059 | 0.191 | 0.307 | Linear |  |  | 0.759 |
| Dose0.5:IntakeRatingLow | 0.483 | 0.188 | 2.576 | Linear |  |  | 0.010 |
| Dose1:IntakeRatingLow | 0.543 | 0.187 | 2.900 | Linear |  |  | 0.004 |
| Dose2:IntakeRatingLow | 0.423 | 0.212 | 1.996 | Linear |  |  | 0.046 |
| Dose5:IntakeRatingLow | 1.230 | 0.480 | 2.564 | Linear |  |  | 0.010 |
| Dose0.5:SexM | -0.082 | 0.188 | -0.440 | Linear |  |  | 0.660 |
| Dose1:SexM | 0.053 | 0.187 | 0.282 | Linear |  |  | 0.778 |
| Dose2:SexM | 0.467 | 0.215 | 2.169 | Linear |  |  | 0.030 |
| Dose5:SexM | -0.411 | 0.457 | -0.899 | Linear |  |  | 0.369 |
| s(Mouse) |  |  |  | Random Effect | 13.567 | 75.988 | 0.000 |
| **Model Statistics** |  |  |  |  |  |  |  |
| BIC | 700.549 |  |  |  |  |  |  |
| R² (Deviance Explained) | 0.875 |  |  |  |  |  |  |
| Random Effect Variance (Mouse) | 0.299 |  |  |  |  |  |  |
| * on log scale |  |  |  |  |  |  |  |

| **Supplemental Table 3**  **Summary of GAM Results for Amphetamine Treatment Self-Administration Fixed Ratio 3 Home Cage Trials** | | | | | | | |
| --- | --- | --- | --- | --- | --- | --- | --- |
| **Effect** | **Estimate*** | **Std. Error*** | **t-value** | **Type Of Term** | **EDF** | **F-value** | **p-value** |
| (Intercept) | 1.423 | 0.054 | 26.261 | Linear |  |  | 0.000 |
| Dose0.5 | -0.412 | 0.074 | -5.603 | Linear |  |  | 0.000 |
| Dose1 | -0.620 | 0.099 | -6.255 | Linear |  |  | 0.000 |
| Dose2 | -1.376 | 0.161 | -8.557 | Linear |  |  | 0.000 |
| Dose5 | -2.126 | 0.290 | -7.332 | Linear |  |  | 0.000 |
| SexM | 0.120 | 0.053 | 2.262 | Linear |  |  | 0.024 |
| s(Time):Dose0 |  |  |  | Thin‐plate spline | 4.235 | 23.071 | 0.000 |
| s(Time):Dose0.5 |  |  |  | Thin‐plate spline | 4.386 | 69.891 | 0.000 |
| s(Time):Dose1 |  |  |  | Thin‐plate spline | 4.757 | 90.674 | 0.000 |
| s(Time):Dose2 |  |  |  | Thin‐plate spline | 3.964 | 93.076 | 0.000 |
| s(Time):Dose5 |  |  |  | Thin‐plate spline | 3.347 | 77.200 | 0.000 |
| s(Mouse) |  |  |  | Random Effect | 4.303 | 6.846 | 0.069 |
| **Model Statistics** |  |  |  |  |  |  |  |
| BIC | 2365.445 |  |  |  |  |  |  |
| R² (Deviance Explained) | 0.632 |  |  |  |  |  |  |
| Random Effect Variance (Mouse) | 0.058 |  |  |  |  |  |  |
| * on log scale  **Supplemental Table 4** |  |  |  |  |  |  |  |
| **Summary of GAM Results for Amphetamine Treatment Free Access Sucrose Home Cage Trials** | | | | | | |  |
| **Effect** | **Estimate*** | **Std. Error*** | **t-value** | **Type Of Term** | **EDF** | **F-value** | **p-value** |
| (Intercept) | 1.954 | 0.084 | 23.366 | Linear |  |  | 0.000 |
| Dose0.5 | -0.234 | 0.076 | -3.091 | Linear |  |  | 0.002 |
| Dose1 | -0.504 | 0.089 | -5.688 | Linear |  |  | 0.000 |
| Dose2 | -0.688 | 0.093 | -7.390 | Linear |  |  | 0.000 |
| Dose5 | -1.276 | 0.125 | -10.244 | Linear |  |  | 0.000 |
| SexM | -0.060 | 0.094 | -0.641 | Linear |  |  | 0.521 |
| s(Time):Dose0 |  |  |  | Thin‐plate spline | 0.985 | 1.815 | 0.153 |
| s(Time):Dose0.5 |  |  |  | Thin‐plate spline | 4.607 | 43.930 | 0.000 |
| s(Time):Dose1 |  |  |  | Thin‐plate spline | 4.806 | 66.980 | 0.000 |
| s(Time):Dose2 |  |  |  | Thin‐plate spline | 3.786 | 87.636 | 0.000 |
| s(Time):Dose5 |  |  |  | Thin‐plate spline | 4.354 | 83.675 | 0.000 |
| s(Mouse) |  |  |  | Random Effect | 7.013 | 28.565 | 0.000 |
| **Model Statistics** |  |  |  |  |  |  |  |
| BIC | 2380.171 |  |  |  |  |  |  |
| R² (Deviance Explained) | 0.527 |  |  |  |  |  |  |
| Random Effect Variance (Mouse) | 0.134 |  |  |  |  |  |  |
| * on log scale  **Supplemental Table 5** |  |  |  |  |  |  |  |
| **Summary of GAM Results for Nicotine Treatment Self-Administration Fixed Ratio 3 Home Cage Trials** | | | | | | | |
| **Effect** | **Estimate*** | **Std. Error*** | **t-value** | **Type Of Term** | **EDF** | **F-value** | **p-value** |
| (Intercept) | 1.390 | 0.081 | 17.079 | Linear |  |  | 0.000 |
| Dose0.5 | 0.194 | 0.086 | 2.250 | Linear |  |  | 0.024 |
| Dose1 | 0.003 | 0.092 | 0.031 | Linear |  |  | 0.975 |
| Dose2 | -0.138 | 0.097 | -1.425 | Linear |  |  | 0.154 |
| Dose5 | -1.183 | 0.212 | -5.583 | Linear |  |  | 0.000 |
| SexM | 0.121 | 0.079 | 1.535 | Linear |  |  | 0.125 |
| s(Time):Dose0 |  |  |  | Thin‐plate spline | 2.272 | 19.977 | 0.000 |
| s(Time):Dose0.5 |  |  |  | Thin‐plate spline | 1.570 | 11.587 | 0.001 |
| s(Time):Dose1 |  |  |  | Thin‐plate spline | 1.952 | 15.742 | 0.000 |
| s(Time):Dose2 |  |  |  | Thin‐plate spline | 1.863 | 33.991 | 0.000 |
| s(Time):Dose5 |  |  |  | Thin‐plate spline | 2.496 | 56.465 | 0.000 |
| s(Mouse) |  |  |  | Random Effect | 6.431 | 14.439 | 0.005 |
| **Model Statistics** |  |  |  |  |  |  |  |
| BIC | 1298.579 |  |  |  |  |  |  |
| R² (Deviance Explained) | 0.527 |  |  |  |  |  |  |
| Random Effect Variance (Mouse) | 0.106 |  |  |  |  |  |  |
| * on log scale  **Supplemental Table 6** |  |  |  |  |  |  |  |
| **Summary of GAM Results for Amphetamine Treatment Free Access Sucrose Home Cage Trials** | | | | | | |  |
| **Effect** | **Estimate*** | **Std. Error*** | **t-value** | **Type Of Term** | **EDF** | **F-value** | **p-value** |
| (Intercept) | 1.491 | 0.144 | 10.337 | Linear |  |  | 0.000 |
| Dose0.5 | 0.150 | 0.187 | 0.802 | Linear |  |  | 0.422 |
| Dose1 | -0.421 | 0.206 | -2.041 | Linear |  |  | 0.041 |
| Dose2 | 0.420 | 0.181 | 2.316 | Linear |  |  | 0.021 |
| Dose5 | -0.397 | 0.211 | -1.877 | Linear |  |  | 0.061 |
| SexM | -0.095 | 0.190 | -0.502 | Linear |  |  | 0.616 |
| Dose0.5:SexM | 0.200 | 0.244 | 0.819 | Linear |  |  | 0.413 |
| Dose1:SexM | 0.509 | 0.259 | 1.961 | Linear |  |  | 0.050 |
| Dose2:SexM | -0.268 | 0.242 | -1.107 | Linear |  |  | 0.268 |
| Dose5:SexM | -0.114 | 0.265 | -0.430 | Linear |  |  | 0.667 |
| s(Time):Dose0 |  |  |  | Thin‐plate spline | 0.879 | 7.014 | 0.005 |
| s(Time):Dose0.5 |  |  |  | Thin‐plate spline | 0.951 | 1.929 | 0.129 |
| s(Time):Dose1 |  |  |  | Thin‐plate spline | 2.140 | 30.532 | 0.000 |
| s(Time):Dose2 |  |  |  | Thin‐plate spline | 1.375 | 30.842 | 0.000 |
| s(Time):Dose5 |  |  |  | Thin‐plate spline | 2.577 | 32.789 | 0.000 |
| s(Mouse) |  |  |  | Random Effect | 4.130 | 8.460 | 0.025 |
| **Model Statistics** |  |  |  |  |  |  |  |
| BIC | 1243.714 |  |  |  |  |  |  |
| R² (Deviance Explained) | 0.411 |  |  |  |  |  |  |
| Random Effect Variance (Mouse) | 0.113 |  |  |  |  |  |  |

* on log scale
